# Low-dose colchicine improves vasorelaxation, reduces arterial remodeling and attenuates blood pressure increases in spontaneously hypertensive rats

**DOI:** 10.1101/2024.07.31.604256

**Authors:** Samuel N. Baldwin, Joakim A. Bastrup, Jennifer van der Horst, Antonella M. G. Formento, Magdalena Dubik, Olga S. Kudryavtseva, Arnela Saljic, Salomé Rognant, Johs Dannesboe, Anthony M. Mozzicato, Thomas Jespersen, Jesper B. Moeller, Jean-Claude Tardif, Morten B. Thomsen, Thomas Jepps

## Abstract

Colchicine, a microtubule depolymerizing agent, is an effective therapy for the secondary prevention of cardiovascular disease and has been approved recently as a novel treatment for atherosclerosis associated with coronary artery disease. Hypertension is a leading cause of cardiovascular disease, yet the impact of colchicine on hypertension has not been studied. We hypothesized that low-dose colchicine could be used to treat hypertension to reduce cardiovascular disease risk. The aim of this study was to administer daily, low dose (0.05 mg/kg/day) oral colchicine for 4 weeks to spontaneously hypertensive rats (SHR) and normotensive controls (WKY) and determine the effect on blood pressure, vascular reactivity, remodeling and inflammation, and left ventricular hypertrophy. Daily blood pressure measurements recorded by telemetry in conscious rats showed colchicine prevented increases in mean arterial pressure observed in the SHRs receiving vehicle over the 4-week treatment period. After the 4-weeks of treatment, 3^rd^ order mesenteric artery vasorelaxations to isoproterenol, sodium nitroprusside and the Kv7.2-5 channel activator, ML213, were enhanced in the SHRs receiving colchicine compared to vehicle. The improved isoproterenol-mediated relaxation was also observed in WKY rats receiving colchicine, and in both the SHR and WKY, this improved effect was attenuated by the β2 adrenoceptor antagonist, ICI118,551. Proteomic analysis of the mesenteric arteries by mass spectrometry revealed that colchicine treatment prevented changes observed when comparing the SHR vehicle group with the WKY vehicle group in proteins associated with extracellular matrix pathways. Immunostaining of 3^rd^ order mesenteric arteries with Sirius red found that colchicine treatment attenuated the increased media thickness of the artery wall observed in SHRs receiving vehicle. Multiplex immunoassay and Western blots revealed colchicine reduced certain inflammatory mediators in the wall of the SHR mesenteric arteries, particularly the nucleotide-binding domain and leucine-rich repeat pyrin containing protein-3 (NLRP3), IL-18, CXCL10, and CXCL2, as well as reducing phosphorylated STAT3. Finally, in the left ventricle of the SHR, colchicine treatment attenuated a number of inflammatory mediators, including NLRP3, IL-1β, and IL-18, and reduced fibrosis and cell size, which are indicative of left ventricular hypertrophy. Overall, we show colchicine has the potential to elicit cardiovascular protective effects in hypertension by targeting multiple cell types.

## Introduction

Colchicine, derived from the autumn crocus plant (*Colchicum autumnale*), is a medication that has been used for millennia, with its first recorded therapeutic use being described >3500 years ago (Dasgeb et al., 2018). Its mechanism of action involves disrupting the process of microtubule formation, which plays a crucial role in various cellular functions, including cell division and intracellular transport. In the absence of colchicine, microtubule polymerization occurs as tubulin dimers are added to the growing end of the microtubule, extending its length. Conversely, during depolymerization, tubulin dimers are removed from the microtubule, shortening its length. This process is termed dynamic instability. Colchicine binds to the β tubulin dimer subunit, interfering with the ability of tubulin dimers to add to the growing ends of microtubules. This disruption inhibits the elongation of microtubules, preventing their normal polymerization, and changing the dynamic instability of the network. This disruption is particularly relevant in immune cells, with colchicine’s anti-inflammatory actions being attributed to suppression of neutrophil chemotaxis (Caner, 1965) and inhibition of the assembly and activation of the nucleotide-binding domain and leucine-rich repeat pyrin containing protein-3 (NLRP3) inflammasome (Martinon et al., 2006). As such, colchicine has been used primarily as an anti-inflammatory in the treatment of gout, familial Mediterranean fever, and acute pericarditis.

Recently, the LoDoCo2 (Low Dose Colchicine) (Nidorf et al., 2020) and COLCOT (Colchicine Cardiovascular Outcomes Trial) (Tardif et al., 2019) clinical trials showed that low-dose colchicine is an effective therapy for secondary cardiovascular prevention. Based on these trials, colchicine has been approved as a repurposed medication to treat patients with coronary artery disease by regulatory authorities like the US Food and Drug Administration.

Hypertension is a major risk factor for cardiovascular disease, including coronary artery disease. In both the LoDoCo2 and COLCOT trials, half of the patients had hypertension. However, no study has investigated the impact of low-dose colchicine on hypertension, nor on the cardiovascular risk specifically associated with hypertension. Colchicine appears to restore arterial relaxation induced by isoproterenol *ex vivo*, which is typically lost in hypertensive rats, by disrupting the microtubule network and the movement of the motor protein dynein (Lindman et al., 2018; van der Horst et al., 2021, 2022). Furthermore, this work was translated clinically in a small cohort of hypertensive humans, which found that a single 0.5 mg dose of colchicine enhanced vascular conductance to isoproterenol and sodium nitroprusside (Ehlers et al., 2023).

Our previous data suggest that colchicine has beneficial effects on arterial reactivity in hypertension. When combined with its known anti-inflammatory properties, we hypothesized that low-dose colchicine could be used to treat hypertension. Thus, we aimed to determine the effect of daily, low-dose oral colchicine treatment for 4 weeks in spontaneously hypertensive rats (SHRs), from proteomic changes in the vascular wall to in vivo changes in blood pressure. Our study confirms, in a rodent model, that colchicine has the potential to improve cardiovascular health in arterial hypertension.

## Methods

### Animals

All experiments were performed in accordance with Directive 2010/63EU on the protection of animals used for scientific purposes and approved by the national ethics committee (approval number 2019-15-0201-00333) and by local Animal Care and Use Committees (institutional approval numbers P21-117 and P24-177). Male normotensive Wistar-Kyoto rats (WKY) and SHRs were purchased from Janvier Labs (France). The rats were group-housed in plastic containers. The rats were maintained at a constant temperature (21-22°C) and humidity (30-35 %) on a 12-h light/dark cycle, with *ad libitum* access to water and rodent chow diet. All animals underwent at least one week of habituation. All *ex vivo* and *in vitro* experiments were performed using 15- to 17-week-old rats euthanized by decapitation while under anaesthesia with isoflurane.

Rats were dosed with either colchicine (0.05 mg/kg, Sigma-Aldrich, Copenhagen, Denmark) dissolved in phosphate buffered saline (PBS) or an equal volume of PBS (vehicle control) via oral gavage, once per day between 9-12 a.m for 4 weeks.

### In vivo blood pressure monitoring

Cohorts of 16 rats, comprised of 8 SHRs and 8 WKYs, were surgically fitted with intra-abdominal aortic radio-telemeters (TRM54P, Kaha Sciences, AD instruments) between 9 and 11 weeks of age. Intra-abdominal aortic radio-telemeters measured blood pressure and temperature continuously. All animals were allowed one week of recovery time after surgery. Rats were placed in their home cages on SmartPads (TR181, Kaha Sciences, AD Instruments), with each SmartPad allowing for recording and charging of one transmitter per cage. Signals were recorded via a PowerLab coupled to LabChart software (AD Instruments, Dunedin, New Zealand). Blood pressure was monitored daily for 1hr in the morning, prior to dosing. Systolic arterial pressure (SAP), diastolic arterial pressure (DAP) and mean arterial pressure (MAP) were measured and calculated via LabChart. Two researchers analyzed the recordings independently.

### Wire myography

Segments of 3^rd^ order mesenteric arteries, ∼2 mm in length, were dissected of fat and peripheral tissue in ice-cold dissecting physiological salt solution containing (in mmol-L^-1^); 120 NaCl, 6 KCl, 1.2 MgCl_2_, 12 glucose, 10 HEPES, with pH balanced at 7.4 via NaOH. Arteries were mounted on 40 μm steel wire within a wire myograph chamber (Danish Myo Technology, Aarhus, Denmark) filled with physiological salt solution (PSS, 5 mL), containing (in mmol-L^-1^); 121 NaCl, 2.8 KCl, 1.6 CaCl_2_, 25 NaHCO_3_, 1.2 KH_2_HPO_4_, 1.2 MgSO_4_, 0.03 EDTA, and 5.5 glucose oxygenated with 95 % O_2_ and 5 % CO_2_ at 37 °C. Arteries were left to warm for 30 min, then normalized to an internal luminal circumference equal to that seen at transluminal pressure of 100 mmHg (13.3 kPa) (Mulvany & Halpern, 1977). Subsequently, vessels underwent an equilibration protocol, whereby they were left for 10 min, then contracted with a high K^+^ physiological salt solution (KPSS) of the following composition (in mmol-L^-1^): 63.5 NaCl, 60 KCl, 1.6 CaCl_2_, 25 NaHCO_3_, 1.2 KH_2_PO_4_, 1.2 MgSO_4_, 0.03 EDTA, and 5.5 glucose to assess viability. Constriction in response to KPSS was allowed to plateau before washing the vessel in PSS (≥4x). This step was repeated, after which endothelial cell integrity was determined by pre-constricting the vessel with 10 μmol-L^-1^ α-1 adrenoreceptor agonist methoxamine, then relaxing the vessel by the addition of 10 μmol-L^-1^ acetylcholine mimetic carbachol. A relaxation of >90 % was deemed as endothelial cell positive.

After equilibration, concentration-effect curves were generated in response to activators of the following: α-1 adrenoreceptor (methoxamine 0.01-30 μmol-L^-1^), thromboxane A2 (TxA2, U46619, 0.001-3 μmol-L^-1^), 5-HT_R_ (5-HT, 0.001-3 μmol-L^-1^) and voltage-gated calcium channels (KCl, 20-120 mmol-L^-1^). After each concentration, an increase in tension was allowed to plateau before the addition of the following concentration. All values were normalized to the second KPSS constriction from the equilibration protocol above. After washing vessels in PSS (≥4x), vessels were allowed >10 min to return to base line tension, then pre-constricted with 10 μmol-L^-1^ methoxamine. After stable tone was achieved, concentration-effect curves were generated in half-log steps in response to activators of the following: β-adrenoreceptor (isoproterenol, 0.001-10 μmol-L^-1^), soluble guanylyl cyclase (sodium nitro prusside, SNP, 0.001-3 μmol-L^-1^), K_V_7.2-5 (ML213, 0.01-10 μmol-L^-1^) and the large conductance calcium-activated potassium channel (BK_Ca_, NS11021, 0.01-10 μmol-L^-1^). Alternatively, vessels were pre-incubated in either β2-selective antagonist ICI 118,551 (100 nmol-L^-1^) for 10 min prior to pre-constriction with methoxamine and the generation of concentration effect curves in response to isoproterenol.

### Proteomics

Dissected mesenteric arteries and aortas were homogenized in protein lysis buffer (in mmol-L^-1^: 50 Tris pH 8.5, 5 EDTA pH 8.0, 150 NaCl, 10 KCl, 1% NP-40, and 1×complete protease inhibitor cocktail (Roche)), as previously described (Bastrup et al., 2022). Supernatants were collected after centrifugation at 11,000 g for 10 min at 4 °C, and protein contents were determined by bicinchoninic acid assay (Thermo Fisher Scientific). Samples were prepared by filter-aided sample preparation (Wiśniewski et al., 2009) using flat spin filters (Microcon-10 kDa), as previously described (Bastrup & Jepps, 2023). In brief, 50 µg protein per sample was diluted in digestion buffer (0.5% sodium deoxycholate in 50 mmol-L^-1^ triethylammonium bicarbonate), and heat-treated for 5 min at 95 °C. Samples were reduced and alkylated in a digestion buffer containing 1:50 (v:v) tris(2-carboxyethyl)phosphine (0.5 mol-L^-1^, Sigma) and 1:10 (v:v) 2-chloroacetamide (0.5 mol-L^-1^, Sigma) for 30 min at 37 °C. Samples were digested overnight at 37 °C with 1 μg Trypsin/LysC mix (Promega). Spin filters were centrifuged at 14, 000 g for 15 min in between incubations. Peptides were desalted using stage-tips containing a poly-styrene-divinylbenzene copolymer modified with sulfonic acid groups (SDB-RPS; 3M) material, as described previously (Bastrup & Jepps, 2023). Peptides were eluted using 80% acetonitrile (Sigma)/2% Ammonia (25%, Sigma), and after vacuum centrifugation, samples were resuspended in 2% acetonitrile (Sigma)/0.1% TFA (Sigma).

### Data acquisition by mass spectrometry

Analyses of samples (70 ng/µL, 1 µL injection) were carried out using a Bruker timsTOF Pro mass spectrometer (Bruker Daltonics) in the positive ion mode with a Captivespray ion source on-line connected to a Dionex Ultimate 3000RSnano chromatography system (Thermo Fisher Scientific). Peptides were separated on an Aurora column with captive spray insert (C18 1.6 μM particles, 25 cm, 75 μm inner diameter; IonOptics) at 60 °C. A solvent gradient of buffer A (0.1 % formic acid) and buffer B (99.9% acetonitrile/0.1% formic acid) over 140 min, at a flow rate of 600 nL/min, were used, respectively. The mass spectrometer was operated in DIA PASEF mode with 1.1 s cycle time, TIMS ramp time of 100.0 ms, and high sensitivity enabled. MS scan range was set to 100 to 1700 m/z.

### Protein identification by mass spectrometry

Raw DIA files from mesenteric artery or aorta samples were analyzed by two separate searches with DIA-NN software (v.1.8.1) (https://github.com/vdemichev/diann) (Demichev et al., 2020) using UniProt FASTA database (UP000002494_10116.fasta (21,587 entries) and UP000002494_10116_additional.fasta (9981 entries), August 2020). The options ‘FASTA digest for library-free search/library generation’ and ‘Deep learning-based spectra, RTs, and IMs prediction’ were enabled. The following settings were used: Digestion protease = Trypsin/P; missed cleavage = 1; max number of variable modifications = 0; N-term M excision = True; C carbamidomethylation = True; peptide length range = 7 to 30; precursor charge range = 1 to 4; precursor m/z range = 300 to 1800; fragment ion m/z range = 200 to 1800; Precursor false discovery rate (FDR) (%) = 1%; Use isotopologs = enabled; MBR = enabled; Neural network classifier = single-pass mode; Protein inference = Genes; Quantification strategy = Any LC (high accuracy); Cross-run normalization = RT-dependent; Library generation = Smart profiling; Speed and RAM usage = Optimal results.

### Bioinformatics

Protein quantities from the unique gene lists generated in DIA-NN (*report.unique_genes_matrix.tsv*) were loaded into Perseus (v1.6.14.0) (Tyanova et al., 2016). Mesenteric artery and aorta data were analyzed in parallel. Data was log2 transformed and samples were annotated and filtered by three valid values in each group. Missing values were only imputed for principal component analysis (width =0.4, down shift = 1.8). Two-sided Student’s *t* tests with permutation-based FDR (<0.05) and 250 randomizations, were used to compare the different groups. Hierarchical clustering was created based on z-scored LFQ values and generated by average linkage, preprocessing with k-means, and Euclidean distance. The STRING (v2.0.1; (Szklarczyk et al., 2015)) and ClueGO (v2.5.10; (Bindea et al., 2009)) apps in Cytoscape (v3.9.1; (Shannon et al., 2003)) were used to generate functionally grouped networks and enrichment analysis of significantly regulated proteins (*p*-value<0.05, non-adjusted). For STRING, the analysis used *Rattus norvegicus* as organism and protein interaction score of 0.4. The following ontology reference sets were used: GO Biological Process, GO Molecular Function, and GO Cellular Component. For ClueGO, the analysis used *Rattus norvegicus (10116)* as organism, ‘GO term Fusion’ was not enabled, and ‘only pathways with pV ≤ 0.05’ were enabled. GO Biological Process (from 10.01.2024) was used as database. The sub-enrichment analysis of inflammation-associated proteins was performed by comparing the mesenteric artery data to a curated list based on a GO term ‘Inflammatory response’ search (https://geneontology.org/, downloaded 04.09.2023).

### Western Blot

Immediately after euthanasia, the left ventricle was divided into two parts along the longitudinal axis and either snap-frozen in liquid nitrogen or formalin-fixed. For protein analysis, snap-frozen left ventricular tissue and mesenteric arteries were thawed and homogenized in an extraction buffer (in mmol-L^-1^: 30 Tris ultrapure, 5 EDTA, 30 NaF, 3% SDS, 10% Glycerol, pH 8.8) and treated with Ultra Turrax for 3 intervals of 5 second each, followed by incubation on ice for 10 min. After homogenization, the sample was centrifuged at 10,000 g for 10 min at 4°C and the supernatant was stored at -80 °C. For Western blot analysis, the samples were prepared by mixing the lysate with DTT and loading buffer (NuPAGE^TM^ LDS Sample Buffer (4X), ThermoFisher). The samples, with a concentration of 50 μg of protein, were subjected to electrophoresis on 10% or 15% acrylamide gels, followed by transfer onto nitrocellulose membranes. Primary antibodies against the following target proteins were then used to probe for NLRP3 (rabbit; 1:1000; NBP1-77080, Novus Biologicals), ASC (mouse; 1:500; sc-514414, Santa Cruz), Caspase-1 (rabbit; 1:500; D7F10, Cell Signaling), IL-1β (rabbit; 1:250; ab9722, Abcam), IL-18 (mouse; 1:250; D046-3, MBL) and GAPDH (rabbit; 1:10,000; EPR16891, Abcam). Protein bands were visualized using fluorescently conjugated secondary antibodies raised in mouse and rabbit IRDye^®^ 800CW (1:20,000; 926-32210, 926-32211, Li-Cor) and rabbit IRDye^®^ 680CW (1:20,000; 926-68071, Li-Cor) and imaged and analyzed on the Odyssey Infrared Imaging System (Li-Cor Biosciences; version 5.2.5).

### Histology and immunohistochemistry of left ventricle

For histological staining, formalin-fixed left ventricular tissue was paraffin-embedded and sectioned at 4 µm before being stained with Sirius red to assess the total amount of ventricular fibrosis. For immunohistochemical staining, formalin-fixed and paraffin-embedded tissue was sectioned at 4 µm and, after deparaffinization, blocking was performed by incubation with 2% (w/v) fraction V bovine serum albumin and 2% (w/v) glycine in PBS at room temperature for 1 hr. Additionally, in order to identify the cell membrane of cardiomyocytes, sections were incubated with Wheat Germ Agglutinin (WGA) diluted in PBS (1:200) (Alexa Flour 594 conjugate, ThermoFisher) for 2 hr at room temperature. Coverslips were mounted with Prolong Gold (ProLong ^TM^ Gold antifade reagent, Invitrogen). Imaging was performed using the Zeiss Axio Scan.Z1, and image analysis was performed using QPath software.

### Histology and Sirius red staining of aorta and mesenteric arteries

Cross sections from mesenteric arteries and aorta were cut at 12 μm thickness using a cryostat microtome (Leica CM3050 S), attached to Superfrost Plus glass slides (VWR), and stored at −80 °C. Tissue sections were stained with Sirius red, as reported previously (Bastrup et al., 2022). Images for aorta were acquired using a ZEISS Axioscan.Z1 slide scanner in brightfield mode with a 20×/0.8 Plan-Apochromat objective lens (ZEISS, Axioscan). Images for mesenteric arteries were acquired using ZEISS Axiovert A1 in brightfield mode with a 63×/1.4 Plan-Apochromat objective lens (ZEISS, Axiovert). Raw .czi files were imported into ZEN 3.2 (blue edition, ZEISS) and the Profile function was used to measure the media thickness.

### Multiplex immunoassay

Dissected mesenteric arteries and aorta were homogenized in PBS by bead beating at 6500 rpm for 4 cycles of 10 seconds. The homogenates were centrifuged at 10,000 g for 10 min at 4 °C, and the supernatants were collected. Levels of C-X-C motif chemokine ligand (CXCL) 10, C-X-C motif chemokine ligand (CXCL) 2, vascular endothelial growth factor (VEGF), interleukin (IL)-17, IL-1α, IL-6, TNFα, IL-4, IL-2, IL-13, IL-10, IL-1β, and C-C motif chemokine ligand 11 (CCL11) were measured using the Procartaplex Rat Custom Panel (13-plex, Thermo Fisher Scientific) following the manufacturer’s instructions. The samples were analyzed using a Bio-plex 200 system (Bio-Rad Laboratories, Inc).

### Statistical Analysis

All data were analysed in GraphPad Prism (version 10.1.2) and presented as mean ± standard error of the mean (SEM). The telemetric blood pressure data was analysed using a 3-way ANOVA followed by a Tukey post-test. In the 3-way ANOVA, the variables were WKY vs SHR, vehicle vs colchicine and weeks. The source of variation, significance from the ANOVA and degrees of freedom are reported. Subsequently, all datasets, unless otherwise stated, were compared using a 2-way ANOVA followed by an uncorrected Fisher’s LSD to ensure only the following, relevant comparisons were made after the 2-way ANOVA: WKY: vehicle vs colchicine; SHR: vehicle vs colchicine; Vehicle: WKY vs SHR; Colchicine: WKY vs SHR. For the concentration effect curves, a 2-way ANOVA followed by a Bonferroni multiple comparisons post-test was used.

## Results

### Low-dose colchicine prevents the increases in blood pressure observed in the SHR over a 4 week treatment period

To determine whether colchicine could affect blood pressure, daily measurements were recorded by telemetry in conscious rats and were averaged over the week. In WKY rats, there was no change in the mean arterial pressure (MAP), systolic arterial pressure (SAP), or diastolic arterial pressure (DAP) in the colchicine or vehicle treated groups across the four weeks (n=3-5; Figure 1).

**Figure 1:**
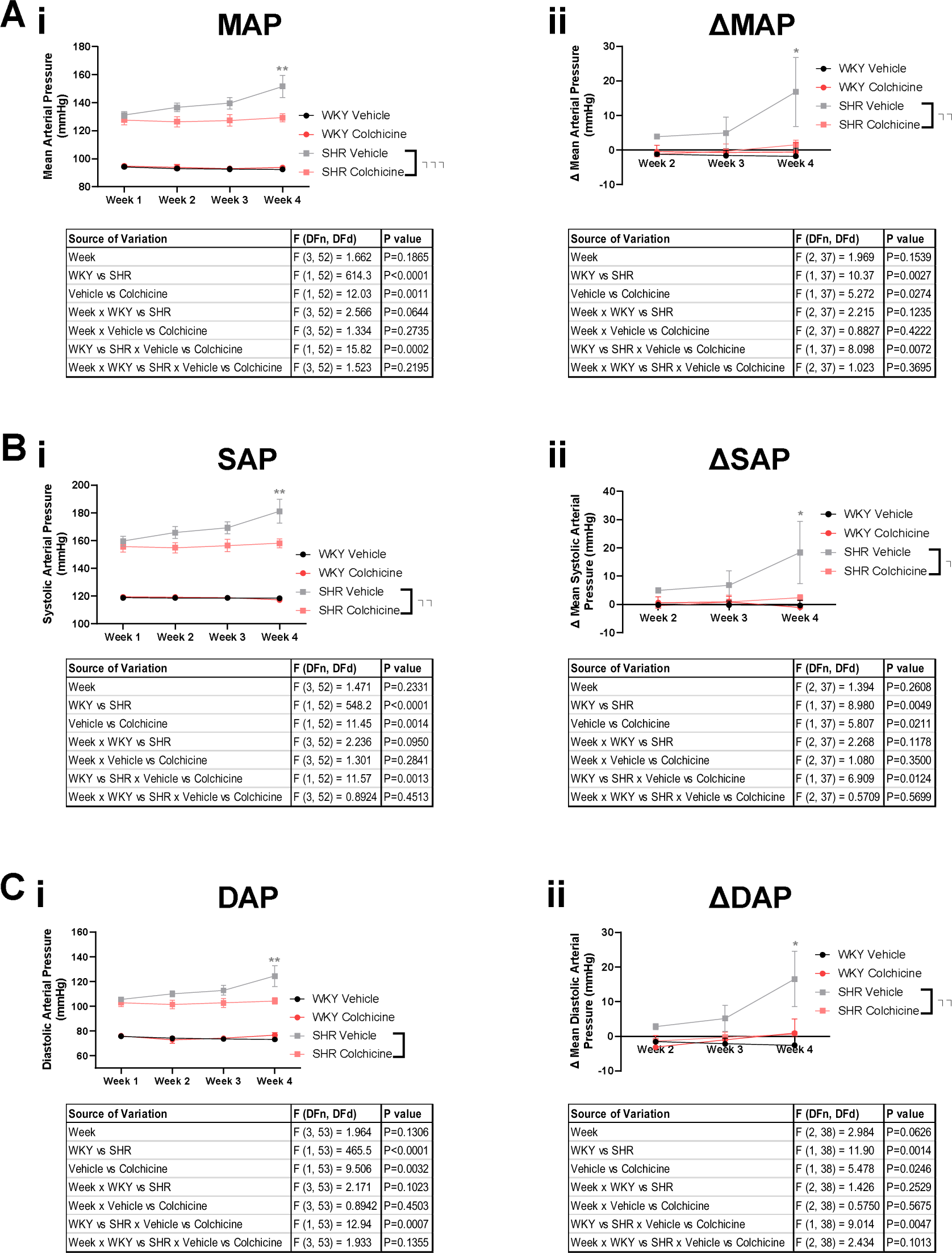
Low-dose colchicine administration to Spontaneously Hypertensive Rats (SHR) and normotensive Wistar Kyoto Rats (WKY) attenuates increases in blood pressure observed in the SHRs over 4 weeks. (A) Mean arterial pressure (MAP), (B) systolic pressure (SAP) and (C) diastolic pressure (DAP) were measured by telemetry and averaged over each week in SHRs and WKY rats (n=3-5). (i) Pressure values and (ii) their mean changes over time are shown relative to week 1. A 3-way ANOVA was performed, and the ANOVA table is presented for each dataset. Statistical significance denoted on the legend corresponds to the WKY vs SHR x Vehicle vs Colchicine comparison. Significance denoted on the graph was determined by a Tukey post-test, * = p<0.05, ** = p<0.01, *** = p<0.001.

In SHRs, MAP, SAP and DAP were significantly higher than in the WKYs (Figure 1; MAP in week 1: 94.2 ± 0.8 mmHg WKY vehicle vs 131.2 ± 2.5 mmHg SHR vehicle; *p<*0.0001; 94.9 ± 1.3 mmHg WKY colchicine vs 127.7 ± 3.5 mmHg SHR colchicine; *p<*0.0001). In the SHR vehicle group, MAP increased over the four weeks to 151.7 ± 8.1 mmHg (Figure 1A; n=3-6; *p=*0.0025), corresponding to a mean increase of 16.8 ± 9 mmHg. These increases were also observed in both the SAP (Figure 1B; 18.4 ± 11 mmHg) and DAP (Figure 1C; 14.9 ± 9.2 mmHg). In contrast, the MAP in the SHR colchicine group did not change from week 1 (127.7 ± 3.5 mmHg) to week 4 (Figure 1A; 129.3 ± 3.0 mmHg; n=5; *p>*0.99). Thus, in week 4, MAP was significantly higher in the SHR vehicle group than the SHR colchicine group (*p=*0.0014). These data show that colchicine prevented the age-dependent increase in blood pressure observed in the SHR vehicle group.

### Chronic colchicine treatment enhances vasorelaxation independent of sensitivity to vasoconstrictors

To investigate the impact of chronic colchicine treatment on resistance artery function, we characterized the effects of a broad spectrum of vasoconstrictors and vasodilators on freshly isolated 3^rd^ order mesenteric arteries.

We observed a significant increase in the potency of β adrenoreceptor- (isoproterenol; Figure 2A-D; *n*=6-9; *p=*0.007 (WKY) and *p=*0.005 (SHR)) and guanylyl cyclase- (SNP; Figure 2B-D; *n*=6-9; *p=*0.01 (WKY) and *p=*0.01 (SHR)) mediated relaxations in both normotensive and hypertensive animals following chronic colchicine treatment. Colchicine also enhanced the K_V_7.2-7.5-mediated (ML213) relaxation in the SHR alone (Figure 2B-D; *n*=7-10; *p=*0.02). Colchicine did not change the EC_50_ of the relaxation evoked by the large-conductance, calcium-activated K^+^ channel (BK_Ca­_) activator, NS1102, although a small enhancement of the relaxation was observed at 3 μmol-L^-1^ in the WKY colchicine group compared to the WKY vehicle group (Figure 2B; n=7-10; *p=*0.03 according to a 2-way ANOVA followed by a Bonferroni post-test). Neither colchicine nor vehicle had any effect on vasoconstriction evoked in response to activators of the α-1 adrenoreceptor (methoxamine 0.01-30 μmol-L^-1^), TxA2_R_ (U46619), 5-HT_R_ (5-HT), nor high K^+^ in hypertensive and normotensive rats (supplemental figure S1; *n*=5-9). Our findings indicate that low-dose colchicine augments physiologically-relevant vasodilatation in arteries from the SHR, independent of a change in the ability of the artery to contract.

**Figure 2.**
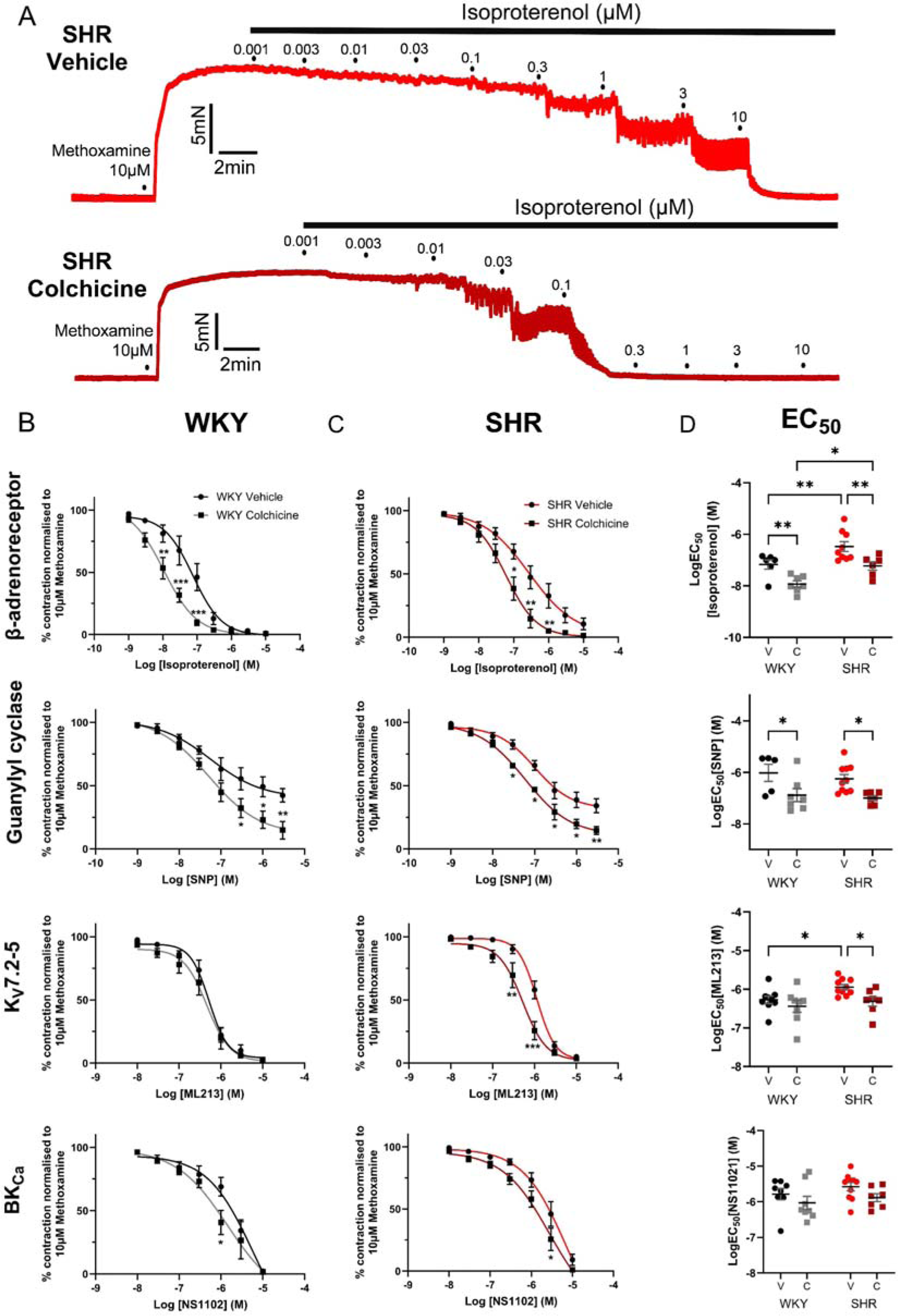
Chronic colchicine treatment enhances vasorelaxation in mesenteric arteries from normotensive (WKY) and hypertensive (SHR) rats. (A) Representative trace of isoproterenol (0.001-10 μmol-L^-1^) mediated vasorelaxation of pre-constricted arterial tone (10 μmol-L^-1^ methoxamine) in 3^rd^ order mesenteric arteries harvested from spontaneously hypertensive rats (SHRs) orally dosed daily for 1 month with vehicle (PBS, red) or 0.05 mg/kg colchicine (burgundy). (B & C) Mean data for relaxation or pre-constricted arterial tone in response to prospective activators of β adrenoreceptor (isoproterenol, 0.001-10 μmol-L^-1^), soluble guanylyl cyclase (sodium nitroprusside, SNP, 0.001-3 μmol-L^-1^), K_V_7.2-5 channels (ML213, 0.01-10 μmol-L^-1^) and BK_Ca_ (NS11021, 0.01-10 μmol-L^-1^) within 3^rd^ order mesenteric arteries harvested from either Wistar Kyoto (n=6-7) or SHRs (n=7-9) dosed with vehicle (WKY black, SHR red) or 0.05 mg/kg/day colchicine (WKY grey, SHR burgundy). Significance was tested with a 2-way ANOVA followed by a Bonferroni post-test and denoted * = p<0.05, ** = p<0.01, *** = p<0.001.

Next, we demonstrate that the isoproterenol-mediated relaxation was attenuated by pre-incubation with the β2 adrenoceptor-selective antagonist ICI 118,551 (100 nmol-L^-1^), in 3^rd^ order mesenteric arteries from vehicle-treated WKY rats (Figure 3A; *n*=6; *p=*0.006). In the WKY rats receiving colchicine, the isoproterenol-mediated relaxation was enhanced (Figure 3A; *n*=5; *p<*0.001), and the effect of ICI 118,551 was greater (Figure 3A; *n*=5; *p<*0.001). In the SHRs receiving vehicle, there was no effect of ICI 118,551 on the isoproterenol-mediated relaxation (Figure 3B; *n=*5; *p=*0.93); however, in the SHR’s receiving colchicine where an enhanced isoproterenol-mediated relaxation was observed (*p=*0.01), ICI 118,551 attenuated the relaxation (Figure 3B; *n*=4-5; *p<*0.001). Collectively, these data, together with our previously published work (van der Horst et al., 2022), demonstrate how low-dose colchicine enhances β adrenoceptor-mediated relaxation in arteries from normotensive and hypertensive rats by increasing the functional contribution of the β2 adrenoceptor.

**Figure 3.**
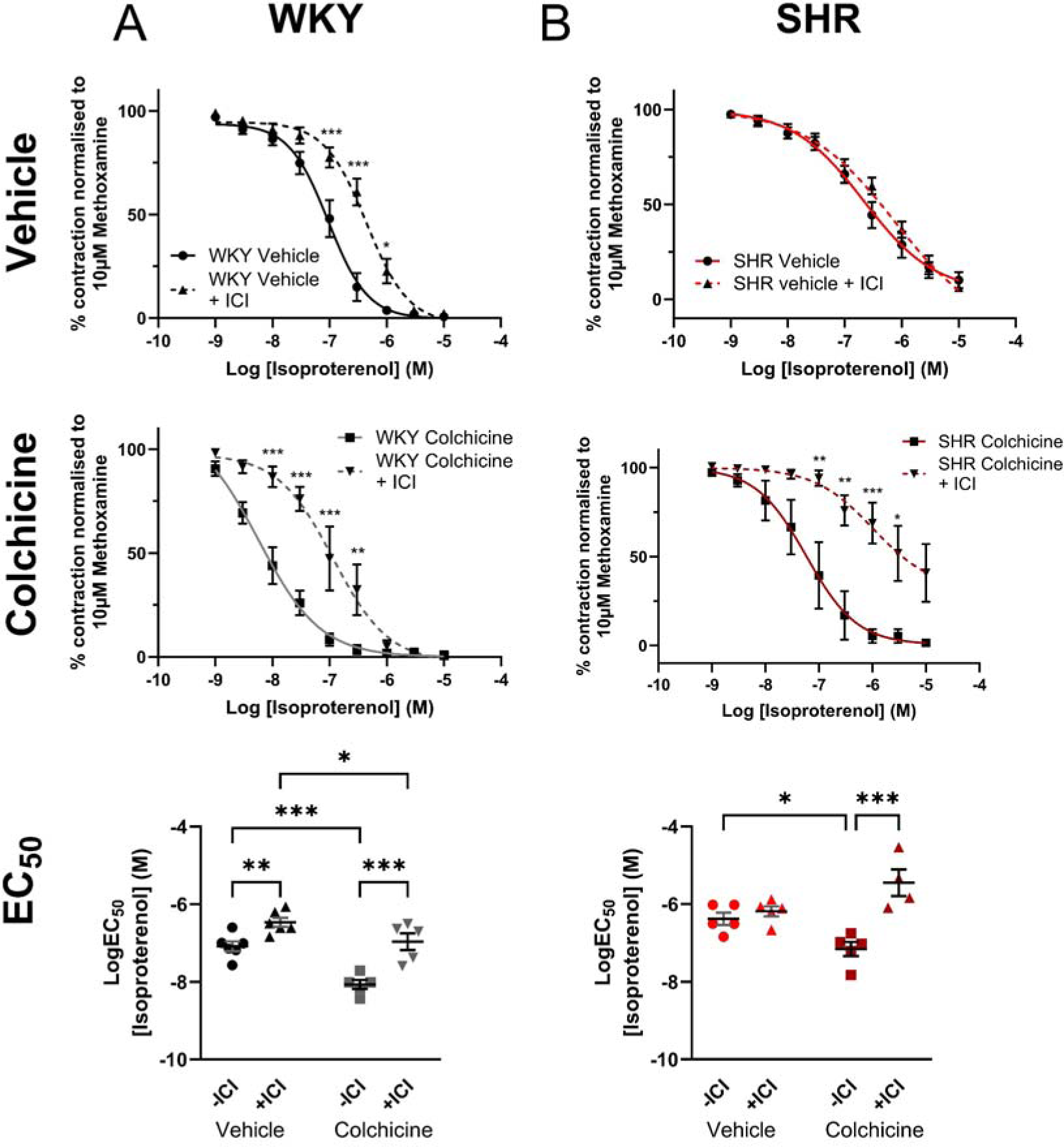
Chronic colchicine treatment restores β2-adrenoreceptor sensitivity in arteries from hypertensive (SHR) rats. Concentration-effect curves showing the response to the β adrenoreceptor activator isoproterenol (0.001-10 μmol-L^-1^) within 3^rd^ order mesenteric arteries harvested from either (A) Wistar Kyoto (WKY, normotensive, n=5-6) or (B) SHRs (n=4-5) dosed with vehicle (WKY black, SHR red) or 0.05 mg/kg/day colchicine (WKY grey, SHR burgundy) in the presence (dashed line) or absence (full line) of ICI-118551 (100 nmol-L^-1^). For the concentration effect curves, statistical significance was tested with a 2-way ANOVA followed by a Bonferroni post-test. EC_50_ values for mean data are displayed in the bottom panel. Here, statistical significance was tested by a 2-way ANOVA followed by an uncorrected Fisher’s LSD, * = p<0.05, ** = p<0.01, *** = p<0.001. (D) EC_50_ values for mean data displayed in (B) & (C). Statistical significance was tested with a 2-way ANOVA followed by a Fisher’s uncorrected LSD (B-D), * = p<0.05, ** = p<0.01, *** = p<0.001.

### Proteomic analysis reveals colchicine reverses changes in the expression of proteins associated with the extracellular matrix in hypertensive (SHR) rats

To characterize molecular changes resulting from low-dose oral colchicine treatment, mass spectrometry analysis was performed on 2^nd^-3^rd^ order mesenteric arteries. The total number of protein IDs were 2776 (Figure 4A). Principal component analysis clearly separated SHRs and WKY controls along component 1, accounting for 12.8% of the variance (Figure 4B). No separation was observed based on colchicine treatment alone. We identified 32 downregulated and 48 upregulated proteins when comparing SHR vehicle to colchicine treatment (Figure 4C). Pathway analysis identified significant associations to ‘angiotensin-activated signaling’ and ‘intermediate filament organization’, among others (Figure 4E & supplementary table S1). We identified 73 downregulated and 75 upregulated proteins when comparing WKY vehicle and WKY colchicine groups (Figure 4D). Pathway analysis highlighted significant associations to ‘cellular detoxification’, ‘positive regulation of DNA biosynthetic process’ and ‘extracellular matrix assembly’ (Figure 4F & supplementary table S2).

**Figure 4:**
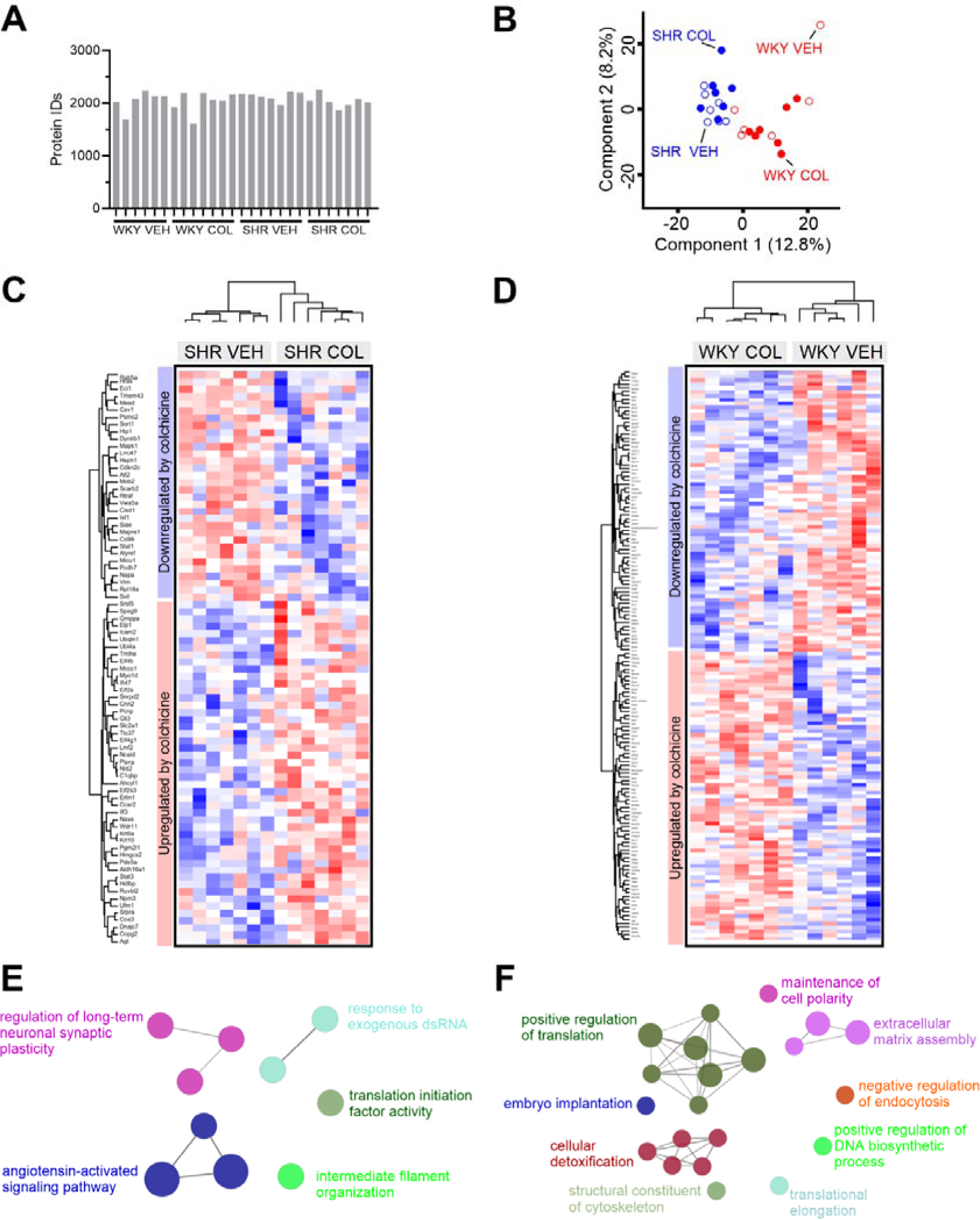
Protein profiling of colchicine treatment in mesenteric arteries. (A) Histogram of unique proteins identified by DIA-MS at 1% false-discovery rate (FDR) identified in spontaneously hypertensive rats (SHR) and normotensive Wistar Kyoto (WKY) controls treated with vehicle (VEH) or colchicine (COL). (B) Principal component analysis (PCA) plot of log2-transformed intensities associated with the samples. Components 1 and 2 are presented. (C & D) Unsupervised hierarchical clustering of significantly regulated proteins identified in spontaneously hypertensive rats (SHR) or normotensive Wistar Kyoto (WKY) controls treated with vehicle (VEH) or colchicine (COL). z-scored values are depicted (range from -3 (blue) to 3 (red)). (E & F) Gene enrichment analysis of differentially expressed proteins (from C and D) using the GO Biological Process database.

To further clarify which proteins were affected by colchicine treatment in mesenteric arteries from hypertensive rats, we compared results from the *WKY vehicle versus SHR vehicle* comparison with the *WKY vehicle versus SHR colchicine* comparison. The majority of significantly regulated proteins were overlapping between both comparisons (=238 proteins). However, 207 proteins were unique to the *WKY vehicle versus SHR vehicle* comparison, while only 146 proteins were unique to the *WKY vehicle versus SHR colchicine* comparison (Figure 5A).

**Figure 5:**
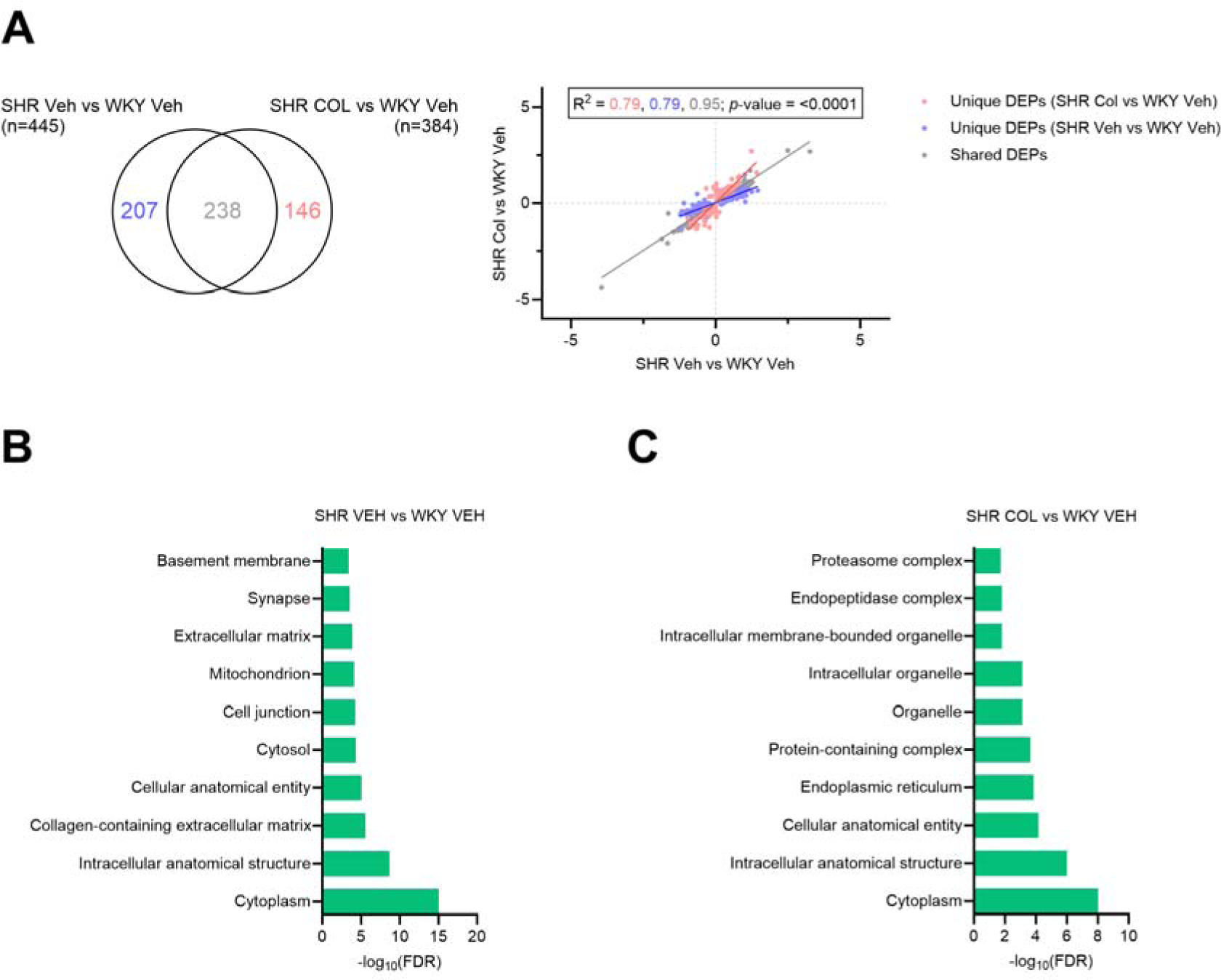
Colchicine treatment attenuates changes in extracellular matrix-associated pathways in hypertensive (SHR) rats. (A) Venn diagram showing unique and shared differentially expressed proteins (DEPs) from two comparisons; Wistar Kyoto (WKY) vehicle (VEH) vs Spontaneously Hypertensive Rat (SHR) vehicle and WKY vehicle vs SHR colchicine (COL). The changes in the expression of proteins in these comparisons is shown in a scatter plot, with R^2^ values calculated. Unique to WKY vehicle vs SHR vehicle = blue, unique to WKY vehicle vs SHR colchicine = red, shared = grey. (B & C) Significance score identified in enrichment analysis of uniquely regulated proteins. Pathway analysis is based on the GO Cellular Component database (top10 pathways based on significance are shown). Log10 transformed significance scores are depicted.

We then visualized the regulation of overlapping and unique proteins identified from both comparisons by plotting their fold change differences. This approach revealed a significant correlation of the overlapping proteins (R^2^ = 0.95), suggesting that these proteins were not affected by treatment. On the contrary, the slope of the unique proteins was different. Here, the differences of unique proteins identified only in the *WKY vehicle versus SHR vehicle* comparison were attenuated and changed towards the baseline by treatment (blue dots in Figure 5A). Similarly, the fold change differences of unique proteins identified in the *WKY vehicle versus SHR colchicine* comparison were changed away from baseline (red dots in Figure 5A).

Pathway analysis identified significant associations to ‘extracellular matrix’ in the blue protein cluster (Figure 5B; supplementary figure S2), suggesting the proteins associated with these pathways are being changed towards what is observed in the WKY vehicle. In the red protein cluster, we mainly found proteins associated to ‘organelles’ as those that are being affected by colchicine (Figure 5C; supplementary figure S2). Overall, these proteomic data suggest that colchicine treatment can prevent changes in the extracellular matrix in SHR mesenteric arteries, which are potentially involved in vascular remodeling.

We also performed proteomic analysis of thoracic aorta segments. The total number of protein IDs was 2941 (supplementary figure S3). Similar to the mesenteric artery data, principal component analysis separated SHRs and WKY controls along component 1 (supplementary figure S3). We identified 33 downregulated and 35 upregulated proteins when comparing SHR vehicle and SHR colchicine groups (supplementary figure S3). However, pathway analysis only identified two significant associations to ‘cellular response to heat’ and ‘lamellipodium morphogenesis’ (supplementary figure S3).

Similar to the mesenteric artery analysis, we also investigated which proteins were affected by colchicine treatment in the aorta segment by focusing on the *WKY vehicle versus SHR vehicle* and *WKY vehicle versus SHR colchicine* comparisons (supplementary figure S4). The majority of significantly regulated proteins were overlapping between both comparisons (=394 proteins). Furthermore, 161 proteins were unique to the *WKY vehicle versus SHR vehicle* comparison, while 177 proteins were unique to the *WKY vehicle versus SHR colchicine* comparison (supplementary figure S4). When plotting the fold change differences from both comparisons, we observed that the unique proteins identified only in the *WKY vehicle versus SHR vehicle* comparison were attenuated and changed towards the baseline by treatment, as observed in the mesenteric arteries. Unlike the mesenteric arteries (Figure 5B), pathway analysis of these proteins did not reveal any significant association to ‘extracellular matrix’, but rather changes in cell cytoplasm and intracellular structure supplementary figure S4).

### Colchicine attenuates remodeling of 3^rd^ order mesenteric arteries from hypertensive (SHR) rats

Because colchicine attenuated changes in proteins associated with several extracellular matrix-related pathways in the SHR, we investigated whether these proteomic changes manifest as changes in the media thickness of 3^rd^ order mesenteric arteries. We stained arterial sections with Sirius red to investigate the thickness of the tunica media layer (Figure 6A). The staining revealed increased media thickness in the SHR-vehicle group compared to the WKY-vehicle group (Figure 6B; *n*=4; *p=*0.0002) and a significantly reduced tunica media thickness in the SHR colchicine-treated group compared to SHR vehicle (Figure 6B; *n*=4; *p=*0.002). Conversely, there was no change in the aorta media thickness when comparing the WKY-vehicle and SHR-vehicle groups, nor was there any effect of colchicine (supplementary figure S5).

**Figure 6:**
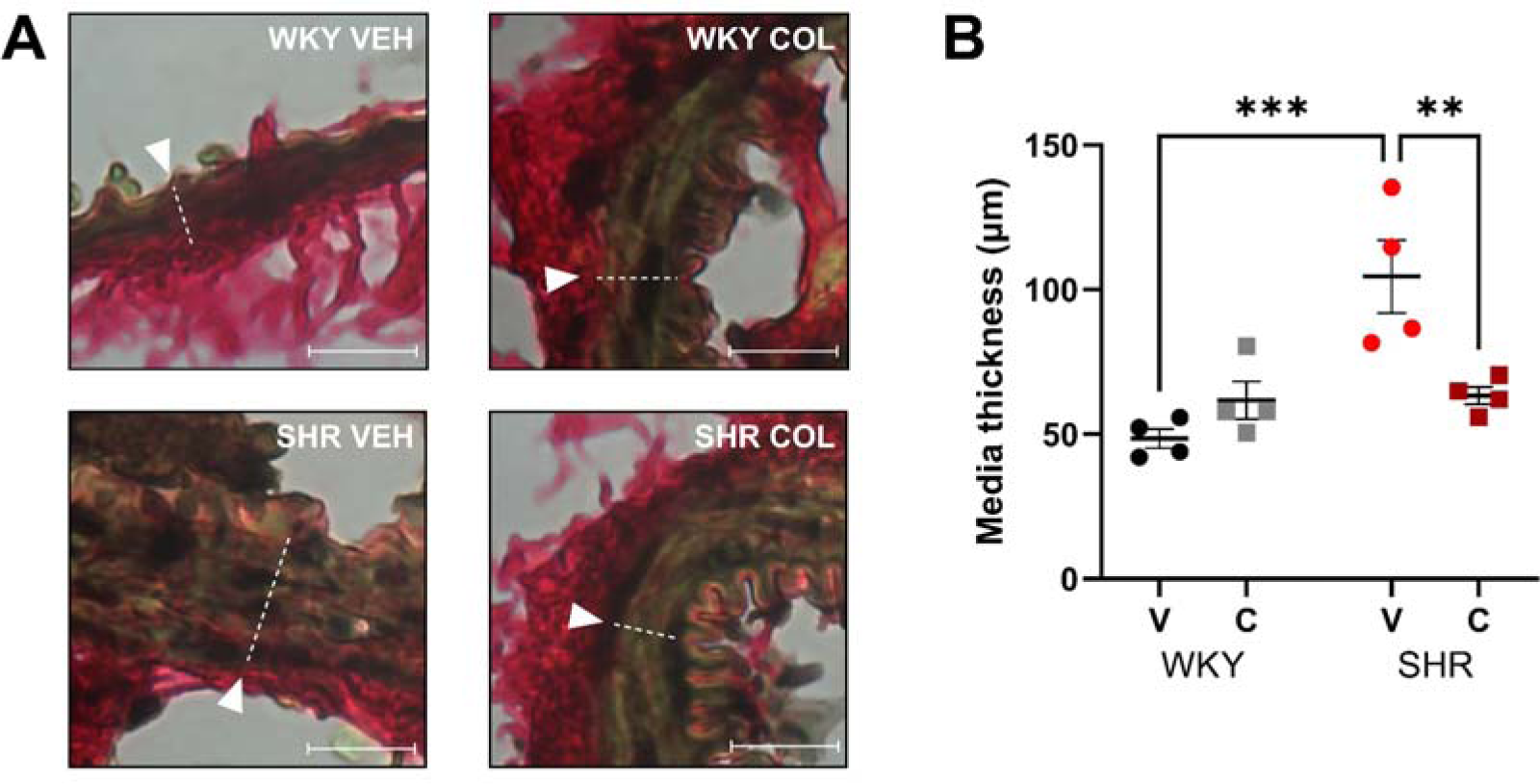
Vascular remodeling of mesenteric artery is reversed by colchicine treatment. (A) Representative Sirius red stained mesenteric arteries from the four groups (Wistar Kyoto (WKY), spontaneously hypertensive rat (SHR), colchicine (COL), vehicle (VEH). White arrow points to line measuring tunica media thickness. Scale bar represents 50 μm. (B) Mean data ± SEM showing the media thickness in WKY and SHR groups receiving vehicle (V) or colchicine (C). (n = 4 in each group; 63 × lens). Statistical significance was determined by a 2-way ANOVA followed by a Fisher’s uncorrected LSD. ** and *** denote significance of p<0.01 and p<0.001, respectively.

### Colchicine reduces CXCL2, CXCL10 and IL-18 expression in the artery wall

Hypertension is associated with low-grade inflammation, which is suggested to contribute to vascular remodeling (Guzik et al., 2024). Although colchicine is fundamentally a microtubule depolymerizing agent, it is considered an anti-inflammatory drug due to its effects on the NLRP3 inflammasome leading to a reduction in the pro-inflammatory cytokine IL-1β, as well as its ability to reduce leukocyte adhesion, proliferation, and chemotaxis. Therefore, we examined whether colchicine had anti-inflammatory effects in the SHR, which might contribute to the reduced media thickness observed in the arterial wall.

Using a multiplex immunoassay platform, we determined the concentration of several inflammatory markers in the mesenteric arteries of the SHR and WKY rats treated with either vehicle or colchicine. Only CXCL10, CXCL2, and VEGF were consistently detected in our samples, whereas TNFα, IL-2, IL-4, IL-10, IL-13, IL-1β and CCL11 were under the detectable range for the assay (see supplementary table 3). Both CXCL10 and CXCL2 were increased in the SHR-vehicle group compared to the WKY-vehicle group (Figure 7A; *n*=3; *p=*0.002 and *p=*0.002, respectively). However, CXCL10 and CXCL2 were reduced in the SHRs treated with colchicine compared to those receiving vehicle (Figure 7A; *n*=3; *p<*0.001 and *p=*0.007, respectively). VEGF was not changed in the SHR vs WKY comparison nor between vehicle and colchicine groups.

**Figure 7:**
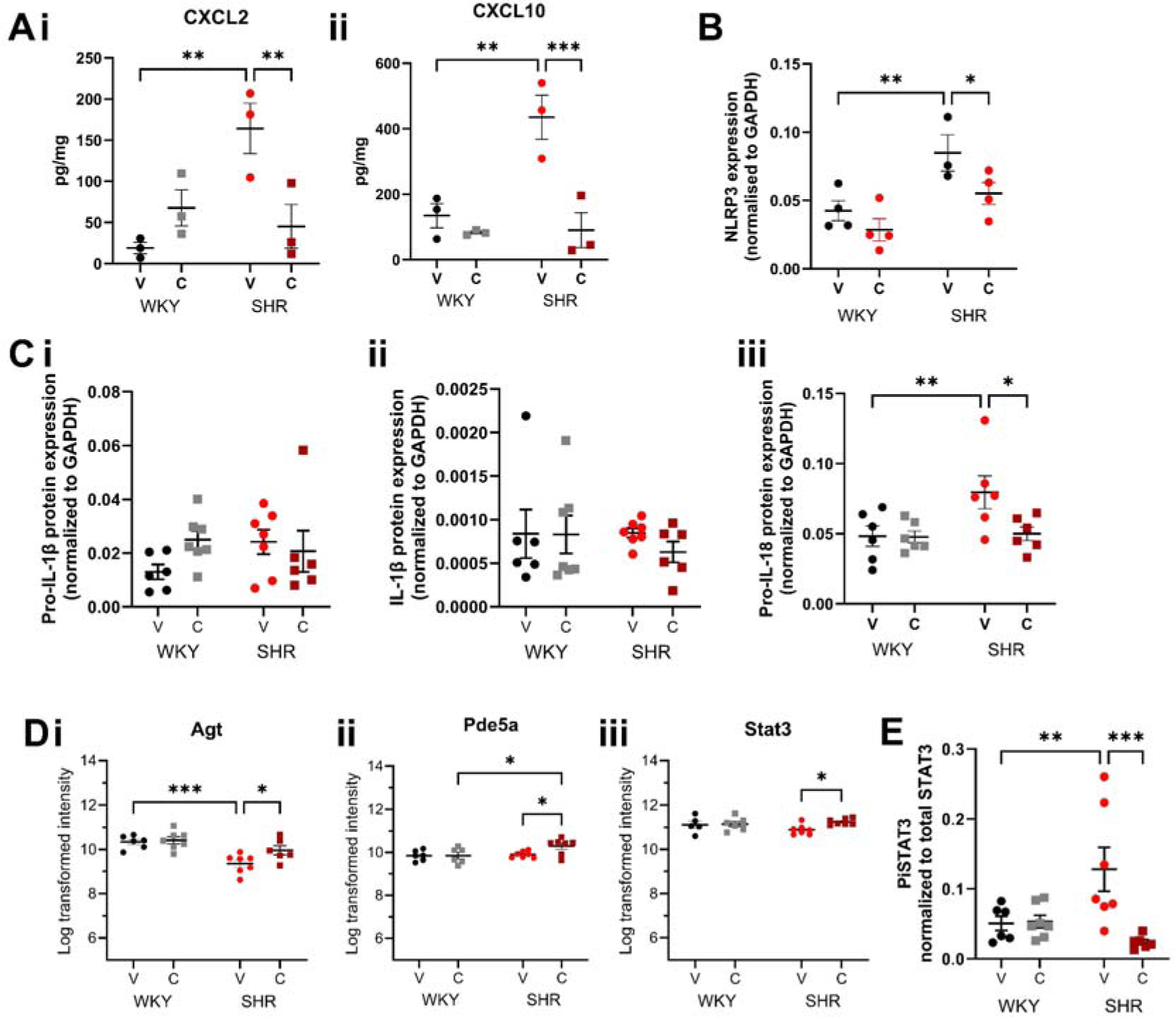
Colchicine treatment reduces the expression of various inflammatory mediators in mesenteric arteries of the SHR. (A) Expression of (i) CXCL2 and (ii) CXCL10 in the mesenteric arteries of Wistar Kyoto (WKY) and Spontaneously Hypertensive Rats (SHR) treated with either vehicle (V) or 0.05mg/kg/day colchicine (C) for 4 weeks. (B) Expression of NLRP3 in in the mesenteric arteries of WKY and SHRs treated with either vehicle (V) or colchicine (C), determined by Western Blot. Full blot can be found in supplementary figure S6. (C) Expression of (i) pro-IL-1β, (ii) cleaved IL-1β, and (iii) pro-IL-18 in the mesenteric arteries of WKY and SHRs treated with either vehicle (V) or colchicine (C), determined by Western Blot. Full blots can be found in supplementary figure S7. (D) From our proteomic data, we identified the expression of (i) Agt, (ii) PDE5A, and (iii) STAT3 to be upregulated in the SHRs receiving colchicine compared to the SHRs receiving vehicle. Here we show the log transformation intensity values of these proteins to determine how expression is changed in the WKY and SHR with vehicle and colchicine treatments. (E) Phosphorylated (activated) STAT3 expression was analysed by Western blot and found to be increased in the SHR-V group compared to the WKY-V and SHR-C groups. Full blots can be found in supplementary figure S9 All statistical significance was determined by a 2-way ANOVA followed by a Fisher’s uncorrected LSD. *, ** and *** denote significance of <0.05, <0.01 and <0.001, respectively.

Previously, colchicine has been associated with an inhibition of the NLRP3 inflammasome and its downstream cytokines, namely IL-1β and IL-18; therefore, we performed Western blots to determine whether their levels were modified in mesenteric arteries of the SHR and whether colchicine treatment affected their expression. In Figure 7B, we show that expression of NLRP3 is increased in the mesenteric arteries of the vehicle-treated SHRs (*n*=3-4; *p=*0.008), which is attenuated in the SHRs receiving colchicine (*n*=3-4; *p=*0.046; supplementary figure S6). We did not detect any change in either pro- or cleaved-IL-1β between the WKY and SHR groups, nor with colchicine treatment (Figure 7C; *n*=6-7; supplementary figure S7). Pro-IL-18 was increased in the SHR-vehicle group compared to the WKY-vehicle rats (Figure 7C; *n*=6-7; *p=*0.009; supplementary figure S7), which was attenuated in the SHRs receiving colchicine (Figure 7C; *n*=6-7; *p=*0.01; supplementary figure S7). We could not reliably detect expression of the activated, cleaved form of IL-18 in these protein lysates (supplementary figure S7).

We then performed a sub-analysis of our proteomics data, enriching these data for inflammation-associated proteins. Out of the total protein background, 136 proteins were present in the inflammation database (supplementary figure S8). Of these, only 3 were significantly upregulated when comparing SHR colchicine to the SHR vehicle group (Stat3, Pde5a, Agt; supplementary figure S6). Further analysis of these proteins showed that Agt is downregulated in the vehicle-treated SHR compared to vehicle-treated WKY, in line with our previous proteomic study on mesenteric arteries from untreated SHRs and WKY rats (Figure 7D; *n*=6-7; *p=*0.0003) (Bastrup et al., 2022). Colchicine increased Agt levels in the SHR to the levels observed in the WKY, thereby accounting for the upregulation we observed (Figure 7D; *n*=6-7; *p=*0.02). Similarly, Stat3 levels were slightly reduced in the vehicle-treated SHR compared to vehicle-treated WKY and were returned to WKY levels in SHRs treated with colchicine (Figure 7D; *n*=6-7; *p=*0.016). Analysis of Stat3 by Western blot found that phosphorylated Stat3 levels were increased in the vehicle-treated SHR (Figure 7E; *n*=6-7; *p=*0.007), but colchicine treatment reduced the phosphorylated Stat3 levels in the SHR to those observed in the WKY (Figure 7E; *n*=6-7; *p<*0.001; supplementary figure S9). Finally, we found that colchicine treatment increased PDE5A levels, only in the SHR (Figure 7D; *n*=6-7; *p=*0.02).

### Colchicine attenuates expression of inflammasome-associated proteins and reduces left ventricular fibrosis in the SHR

Western blot analysis was used to examine the effects of colchicine on the NLRP3 inflammasome (NLRP3, Caspase-1, ASC) and on pro-inflammatory cytokines (IL-1β and IL-18) in the left ventricle of SHR and WKY rats treated with either vehicle or colchicine (n=5-11). We observed a significant difference in the expression levels of NLRP3, pro-Casp-1, Caspase-1, Pro-IL-18, and IL-18 between the WKY and SHR groups receiving vehicle (Figure 8A), indicating an increase in inflammasome activation in the latter. Colchicine treatment in the SHR significantly reduced the levels of NLRP3, Caspase, IL-1β, Pro-IL-18, and IL-18 compared to the SHRs receiving vehicle (Figure 8A). No changes were detected in the protein levels of ASC nor Pro-IL-1β (Figure 8A). All Western blots can be found in supplementary figure S10. Given the correlation between the inflammatory process and pro-fibrotic remodeling of the myocardium, we conducted histological and immunohistochemical analyses, using Sirius Red and WGA staining to investigate fibrosis and cellular hypertrophy, which are indicative of left ventricular hypertrophy. Cell size was increased in SHR vehicle compared to WKY vehicle, indicating cellular hypertrophy in the former group (Figure 8B and 8C). With colchicine treatment, we found a significant reduction in both percentage fibrosis and cell size in SHRs compared to SHRs receiving vehicle.

**Figure 8:**
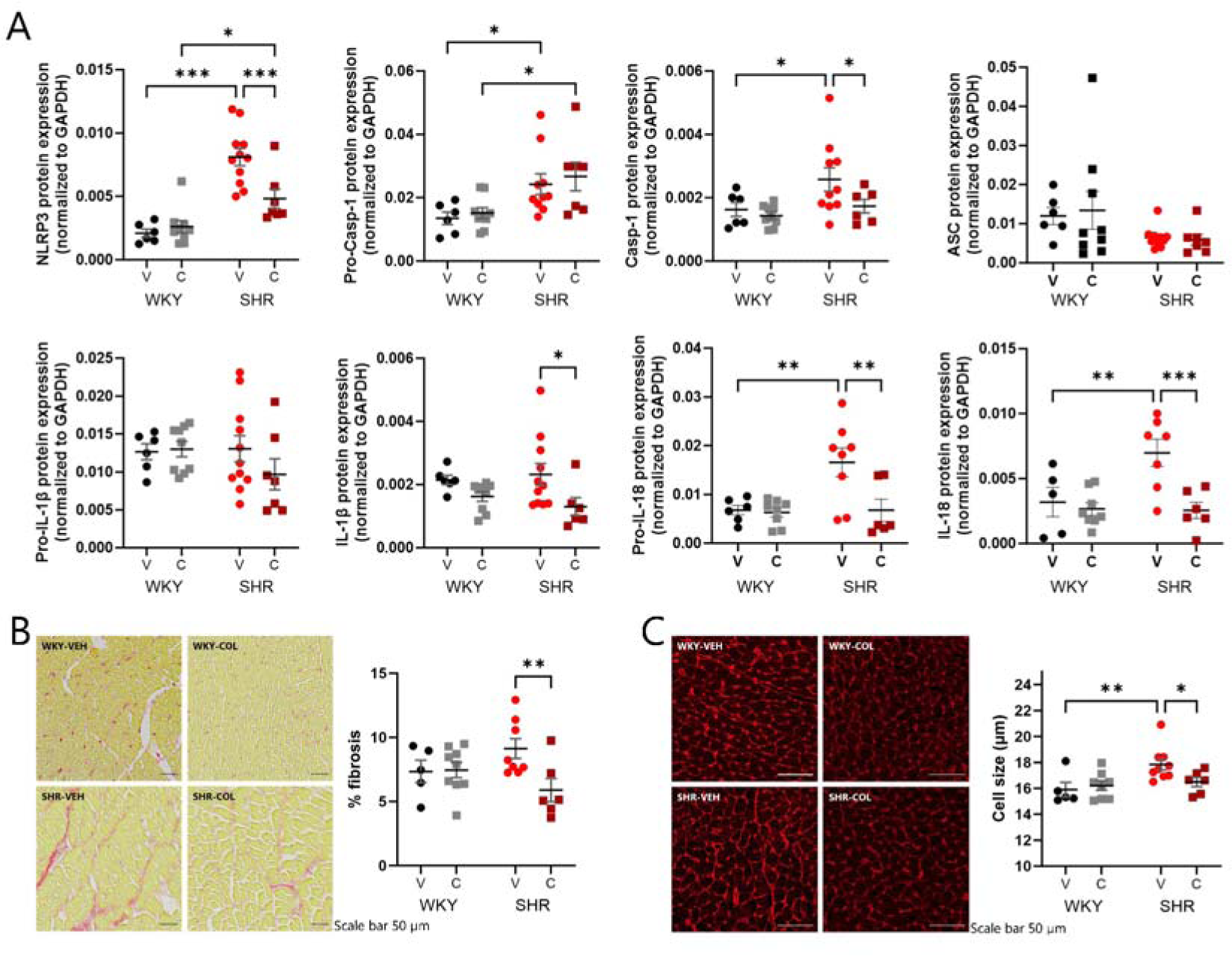
Colchicine treatment reduces the expression of inflammatory mediators, percentage of fibrosis, and cell size in the left ventricle of the SHR. (A) Western Blot analysis showing the expression levels of NLRP3 inflammasome components (NLRP3, Caspase-1, and ASC) and pro-inflammatory cytokines (IL-1β and IL-18) in the left ventricle of WKY and SHRs treated with either vehicle (V) or colchicine (C). Full blots can be found in supplementary figure S10. (B) Quantification of myocardial fibrosis in the left ventricle of WKY and SHRs treated with either vehicle (V) or colchicine (C), determined by histological analysis with Sirius Red staining. (C) Measurement of cell size in the left ventricle of WKY and SHRs treated with either vehicle (V) or colchicine (C), determined by immunohistochemical analysis using WGA staining. All statistical significance was determined by a 2-way ANOVA followed by a Fisher’s uncorrected LSD. *, ** and *** denote significance of <0.05, <0.01 and <0.001, respectively.

## Discussion

As a microtubule-disrupting agent, colchicine has the ability to affect the function of several cell types in the body (Leung et al., 2015). Through its ability to destabilize the microtubule network, colchicine has anti-inflammatory properties that have been recognized for millennia. Recently, colchicine has been approved for the secondary prevention of cardiovascular disease. Although several studies have investigated the cardiovascular protective mechanism of colchicine, often attributing the effect to its anti-inflammatory properties, we do not fully understand how colchicine protects against cardiovascular events and whether some patients might benefit more than others from daily, low-dose colchicine.

In this study, we aimed to investigate the impact of low-dose colchicine in rats with hypertension to ascertain whether some of its reported cardiovascular protective effects might be due to improved vasodilatation, reduced blood pressure, attenuated vascular inflammation and/or effects on vascular and cardiac remodeling. Remarkably, we found that just 4 weeks of colchicine treatment, administered early after the onset of hypertension, was able to improve all these parameters.

The onset of hypertension in the SHR occurs between 6-12 weeks of age, and blood pressure continues to increase steadily thereafter. At 13 weeks, once hypertension was established, we began daily, low-dose, oral administration of colchicine. Although it did not actively lower blood pressure in these rats, colchicine was able to prevent further increases observed in the vehicle-treated SHRs. Thus, colchicine can have an immediate effect on hypertension by limiting the development of the disease. It remains to be determined whether longer-term treatment can sustain this effect and even start to reduce blood pressure. Moreover, future studies will determine whether colchicine can enhance the blood pressure lowering effect of current anti-hypertensive medication.

Vascular smooth muscle cells express both β1 and β2 adrenoreceptors. Evidence suggests that β1 adrenoceptors dominate functionally within mesenteric artery smooth muscle cells (Briones et al., 2005; Graves & Poston, 1993; White et al., 2001), though β2 adrenoceptors contribute. β2 adrenoceptors are internalized in a microtubule/dynein motor-protein dependent process following activation by isoproterenol (Claing, 2002; Jung et al., 2016; Shiina et al., 2000; Suzuki et al., 1992; van der Horst et al., 2022). Previously, we have established that the microtubule network can regulate the movement of certain ion channels and receptors in vascular smooth muscle. We have shown that the Kv7.4 channel, as well as the β2 adrenoceptor, can be trafficked away from the vascular smooth muscle cell membrane along the microtubule network by the motor protein dynein (van der Horst et al., 2021, 2022). *Ex vivo* treatment of arteries with colchicine to disrupt the microtubule network or ciliobrevin to inhibit dynein movement, increased the membrane abundance of Kv7.4 channels and attenuated the agonist-induced internalization of the β2 adrenoceptor, thereby increasing their functional contribution to the isoproterenol-mediated relaxation in both WKY and SHR mesenteric arteries. Whether colchicine could elicit the same effects when administered *in vivo* to conscious rats was not known. This study establishes that oral colchicine treatment enhances responses to isoproterenol in both SHR and WKY mesenteric arteries, together with the relaxation to SNP. Interestingly, the relaxation to the Kv7.2-7.5 activator, ML213, was only enhanced in the SHR, where Kv7 channel function is impaired, as reported in several studies previously (Barrese et al., 2018; Carr et al., 2016; Jepps et al., 2011). These findings establish that *in vivo* colchicine treatment can effectively restore Kv7 channel function in arteries from hypertensive rats and enhance physiologically-relevant relaxations, thereby improving the ability of arteries to dilate. Furthermore, we confirm our previous finding that the enhanced β adrenoceptor-mediated relaxation is elicited by increasing the contribution of the β2 adrenoceptor in vascular smooth muscle, most likely by disrupting the microtubule-dependent, agonist-induced internalization of the receptor.

In early-onset hypertension, resistance arteries undergo remodeling that is associated with an increased media/lumen ratio. In this study, we found that the mesenteric arteries of vehicle-treated SHRs had increased media thickness. In the rats treated with colchicine, the media thickness was reduced significantly, suggesting that colchicine was able to protect against hypertension-induced remodeling since these rats still had hypertension. Hypertension-induced remodeling is a maladaptive process leading to changes in the elastic and mechanical properties of the vascular wall, which compromises blood flow and contributes to increased blood pressure. Preventing such vascular remodeling in hypertension can help to relieve the progression and burden of the disease (Buus et al., 2013).

Remodeling is known to occur due to several inflammatory, apoptotic, and oxidative stress pathways leading to changes in actin polymerization, collagen and elastin levels and activation of matrix metalloproteinases (Castorena-Gonzalez et al., 2014). Our proteomics data suggested that especially the extracellular matrix and mitochondria-associated pathways were affected positively by colchicine treatment. The proteins involved in extracellular matrix changes associated with colchicine in the SHR included Nid2, Lamc1, Mgp, Gsto1, Fgl2, Ccn2, Lama1, Rpsa, Acan, Eln, Fbln1, Abi3bp, Col14a1, Lox, Thbs1, Efemp2 and Lama5. Elastin (Eln) provides elasticity to the arterial wall and was slightly upregulated by colchicine treatment in SHRs. Accumulation of Aggrecan (Acan) has been identified in cardiovascular diseases, such as aortic aneurysm and atherosclerosis (Koch et al., 2020), and in our study colchicine treatment reduced Acan levels in SHRs. Our findings suggest that colchicine has the potential to prevent arterial remodeling; however, the mechanism by which this occurs is not completely clear.

Inflammatory dysregulation is often observed in primary hypertension and is correlated with vascular remodeling. The chronic, low-grade inflammation associated with hypertension is known to affect the kidneys and vascular wall, thereby contributing to the pathogenesis of the disease. This dysregulation is complex, involving several different inflammatory mediators, cytokines, cell types, and various leukocytes (Guzik et al., 2024; Krishnan et al., 2014). In the current study, we found that pro-IL-18 expression was increased in the mesenteric arteries of SHRs receiving vehicle, while colchicine treatment attenuated this increase. Pro-IL-18 is cleaved by the intracellular cysteine protease caspase-1, into active IL-18, which is secreted from the cell. In this process, pro-caspase-1 is first activated into active caspase-1 by the NLRP3 inflammasome. Colchicine is known to inhibit the assembly of the NLRP3 inflammasome (Martinon et al., 2006), as well as suppress its protein expression under pro-inflammatory conditions (Liu et al., 2023). Our data show that colchicine was able to attenuate both NLRP3 and IL-18 expression in SHR mesenteric arteries. IL-18 has been strongly linked with hypertension (Rabkin, 2009). The association between IL-18 and hypertension was highlighted in a meta-analysis that showed a significant positive correlation between blood pressure and circulating IL-18 levels (Rabkin, 2009). Furthermore, IL-18 levels are correlated positively with media thickness (Yamagami et al., 2005). Under certain conditions (including increased angiotensin II and catecholamine release), IL-18 can be released by monocytes, macrophages and vascular endothelial cells. Increased IL-18 promotes vascular smooth muscle cell proliferation and migration, thereby enhancing the remodeling processes associated with hypertension (Chandrasekar et al., 2006; Valente et al., 2012). Interestingly, siRNA-mediated knockdown of IL-18 attenuated angiotensin II-induced vascular smooth muscle cell proliferation in culture, suggesting a critical role for IL-18 in the remodeling promoted by angiotensin II. Both increased angiotensin-II and IL-18 (Kalina et al., 2000) are known to activate (phosphorylate) STAT3, which is associated with the upregulation of genes associated with cardiac remodeling (Han et al., 2018), but whether this pathway is important in vascular remodeling remains to be determined. Our proteomics data revealed an upregulation of STAT3 in SHRs treated with colchicine, and further investigation revealed that this upregulation returned the expression level to that seen in the WKY rats. Moreover, colchicine treatment reduced phosphorylated STAT3 compared to the SHR-vehicle group. Interestingly, colchicine was found recently to inhibit STAT3 phosphorylation in lung tissue of mice with sepsis-induced acute lung injury. This inhibition attenuated NLRP3 activation by repressing its promoter acetylation via the STAT3/EP300 complex (Liu et al., 2023). Further investigations are warranted to determine whether STAT3 activation and NLRP3 play a role in vascular remodeling.

In this study, we also screened mesenteric artery tissue for several cytokines and inflammatory mediators. Most were below the level of detection in the hypertensive rats, suggesting that the degree of inflammation in the vascular wall in a model of early-onset hypertension is low. We did, however, detect changes in CXCL10 and CXCL2 consistent with previous studies linking elevated levels of both chemokines to hypertension (Antonelli et al., 2012; Martynowicz et al., 2014). Interestingly, colchicine reduced the levels of both CXCL10 and CXCL2 in the SHR mesenteric arteries. CXCL2 and CXCL10 are chemokines that stimulate the chemotaxis of neutrophils and CD4+ T-cells, respectively. CXCL10 is secreted by several cell types, including endothelial cells, and promotes chemoattraction of activated T lymphocytes, natural killer cells, and monocytes (Julian et al., 2021). In the context of hypertension, increased CXCL10 leads to increased infiltration of T cells in the kidney, exacerbating hypertension and tissue damage (Antonelli et al., 2012; Youn et al., 2013). Prevention of T-cell migration and activation is known to attenuate hypertension in several animal models of hypertension. CXCL2 expression is increased in the aorta of the SHR and human internal mammary artery of hypertensive patients in comparison with normotensive patients (Jin et al., 2022). Similarly, in mesenteric arteries, we found CXCL2 levels were elevated in the SHR, which was reduced by colchicine. CXCL2 promotes the recruitment of neutrophils, contributes to increased oxidative stress and the infiltration of immune cells, and induces the expression and release of inflammatory mediators such as IL-1β and IL-18 (Boro & Balaji, 2017; Guo et al., 2020). A predominant mechanism of colchicine’s anti-inflammatory effects is through the interference of neutrophil adhesion, recruitment, and chemotaxis, as well as preventing the assembly of the NLRP3 inflammasome. Our data pinpoint several inflammatory mediators targeted by colchicine in the vascular wall of the SHR, leading to a reduction in specific inflammatory pathways that have been associated with hypertension in humans, which may be beneficial in early-onset hypertension as well as in vascular remodeling. Taken together, these findings suggest a novel mechanism by which colchicine may reduce immune cell chemotaxis and vascular inflammation associated with hypertension.

Finally, increased afterload on the heart due to hypertension can induce cardiac fibrosis and remodeling, which are established risk factors in several heart diseases, such as heart failure and arrhythmias (Fuchs & Whelton, 2020; He et al., 2001; Lip et al., 2017). Thus, we wanted to ascertain whether colchicine could protect the heart from cardiac remodeling. Despite colchicine-treated SHRs still having hypertension, we found that colchicine decreased left ventricular fibrosis compared to vehicle control. Cell size analysis showed a significant reduction in cellular hypertrophy in the left ventricle of colchicine-treated SHRs, indicating its ability to reduce hypertrophic changes. Activation of the cardiac inflammasome pathway promotes cardiac remodeling (Scott Jr et al., 2021; Song et al., 2023). We found several components of the NLRP3 inflammasome to be upregulated in SHRs receiving vehicle, including NLRP3, pro-Casp-1, Casp-1, IL-1β, Pro-IL-18, and IL-18. Colchicine treatment attenuated the expression of NLRP3, pro-Casp-1, IL-1β, Pro-IL-18 and IL-18 in the left ventricle, suggesting that colchicine reduced inflammasome-mediated inflammation. This reduction in inflammation may underlie the observed decrease in fibrosis representing a significant defense against cardiac remodeling induced by hypertension. This aligns with previous findings that increased NLRP3 inflammasome activity promotes structural and electrical remodeling of the heart (Li et al., 2023; Yao et al., 2018) and that colchicine can effectively attenuate NLRP3 inflammasome formation and signaling, as well as other inflammatory mediators (Martínez et al., 2018; Opstal et al., 2020).

In summary, our study shows that daily, low-dose colchicine administered for 4 weeks to SHRs improved vasodilatation and reduced vascular remodeling and inflammation in the vascular wall, which may have contributed to the attenuation of blood pressure increases observed in the untreated rats. Colchicine also protected the heart from fibrosis associated with hypertension. Importantly, these cardiovascular protective effects were observed despite the SHR still having hypertension. It remains to be determined whether longer-term treatment can reduce blood pressure and/or enhance the effect of current anti-hypertensive medication. Future clinical trials, such as NCT04916522, will determine whether colchicine has the potential to contribute to the treatment of hypertension.

## Acknowledgements

We would like to thank the Core Facility for Integrated Microscopy and the Telemetry Unit (both University of Copenhagen) for their technical assistance. We are grateful to Karin Larsen (University of Copenhagen) for performing the surgeries to place the telemeters in the rats.

## Data Availability Statement

All original data pertaining to the results of this study will be made available upon reasonable request to the corresponding author.

