## Supplementary Material for "Low-dose colchicine improves vasorelaxation, reduces arterial remodeling and attenuates blood pressure increases in spontaneously hypertensive rats"

***Supplementary Results***

*Figure S1*

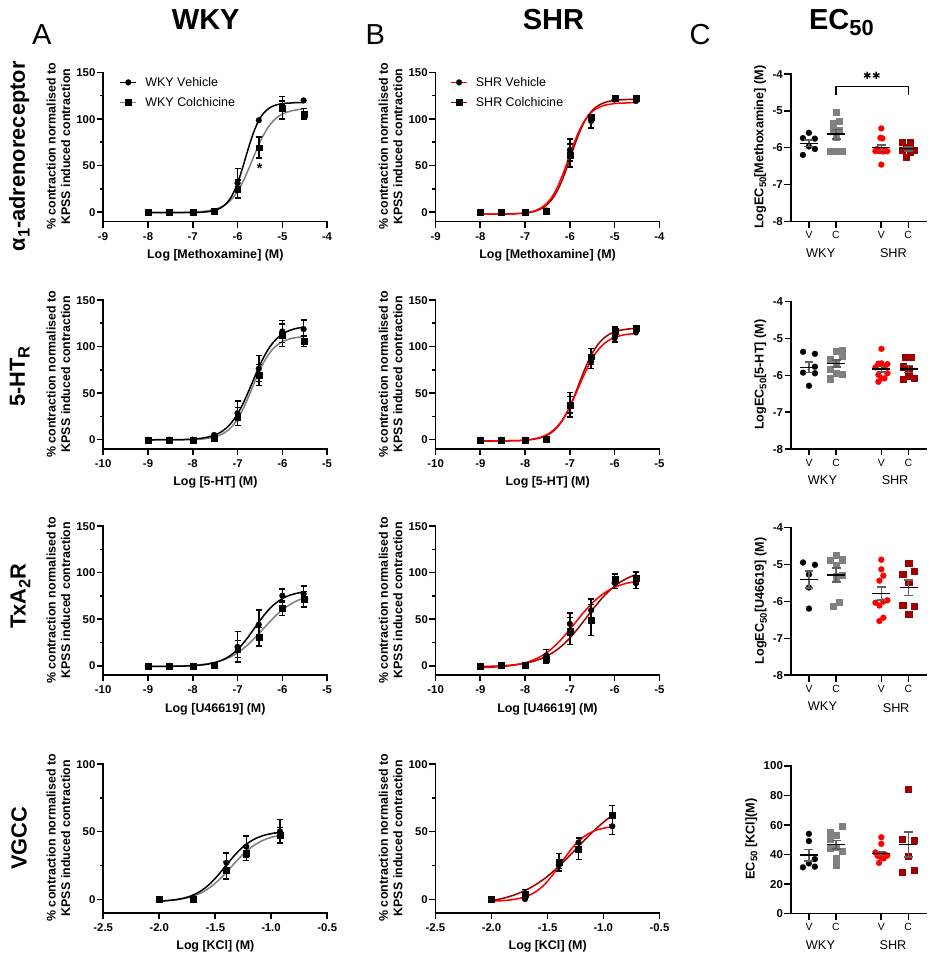

***Supplementary figure 1. Chronic colchicine treatment has no effect on vasoconstriction in arteries from normotensive (WKY) and hypertensive (SHR) rats.***

*Mean data for contraction in response to prospective activators of α-adrenoreceptor (methoxamine, 0.01-10 μmol-L^-1^), 5-HTR (5-HT, 0.001-3 μmol-L^-1^), TxA2R (U46619, 0.001-3 μmol-L^-1^) and voltage gated Ca2+ channels (VGCC, KCl, 10-120 mmol-L^-1^) within 3rd order mesenteric arteries harvested from either (A) Wistar Kyoto (WKY, normotensive, n=5-7) or (B) SHRs (n=7-9) dosed with vehicle (WKY black, SHR red) or 0.05 mg/kg colchicine (WKY grey, SHR burgundy). (C) EC_50_ values for mean data displayed in (A) and (B). Statistical significance was determined by 2-way ANOVA followed by a Fisher’s uncorrected LSD (A-C), **=p<0.01.*

*Figure S2*

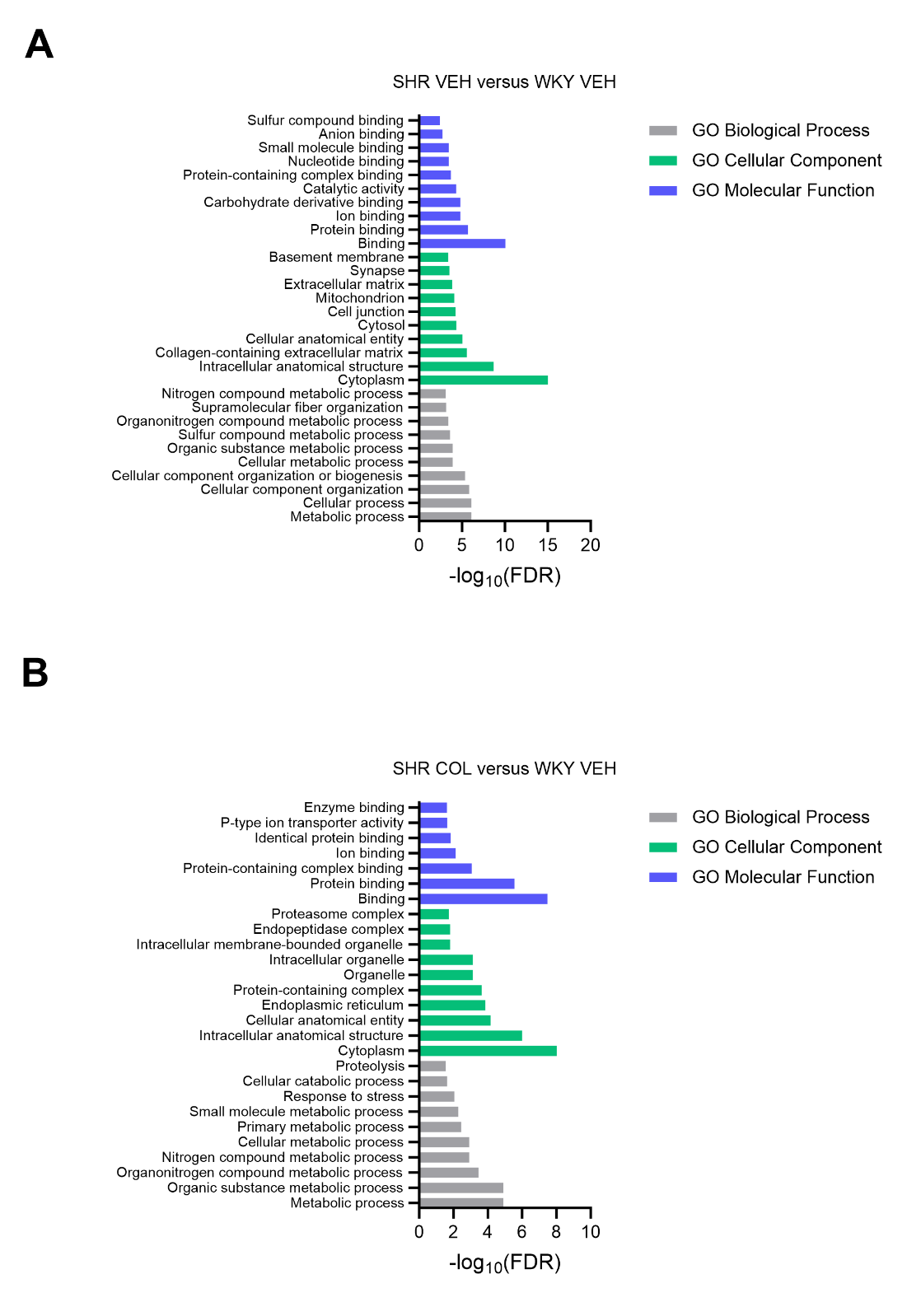

***Supplementary figure S2: Extended pathway analysis of mesenteric artery proteomics data.***

*Significance score identified in enrichment analysis of uniquely regulated proteins. Pathway analysis is based on databases; GO Biological Process,* *GO Cellular Component, and GO Molecular Function. Log10 transformed significance score is depicted.*

*Figure S3*

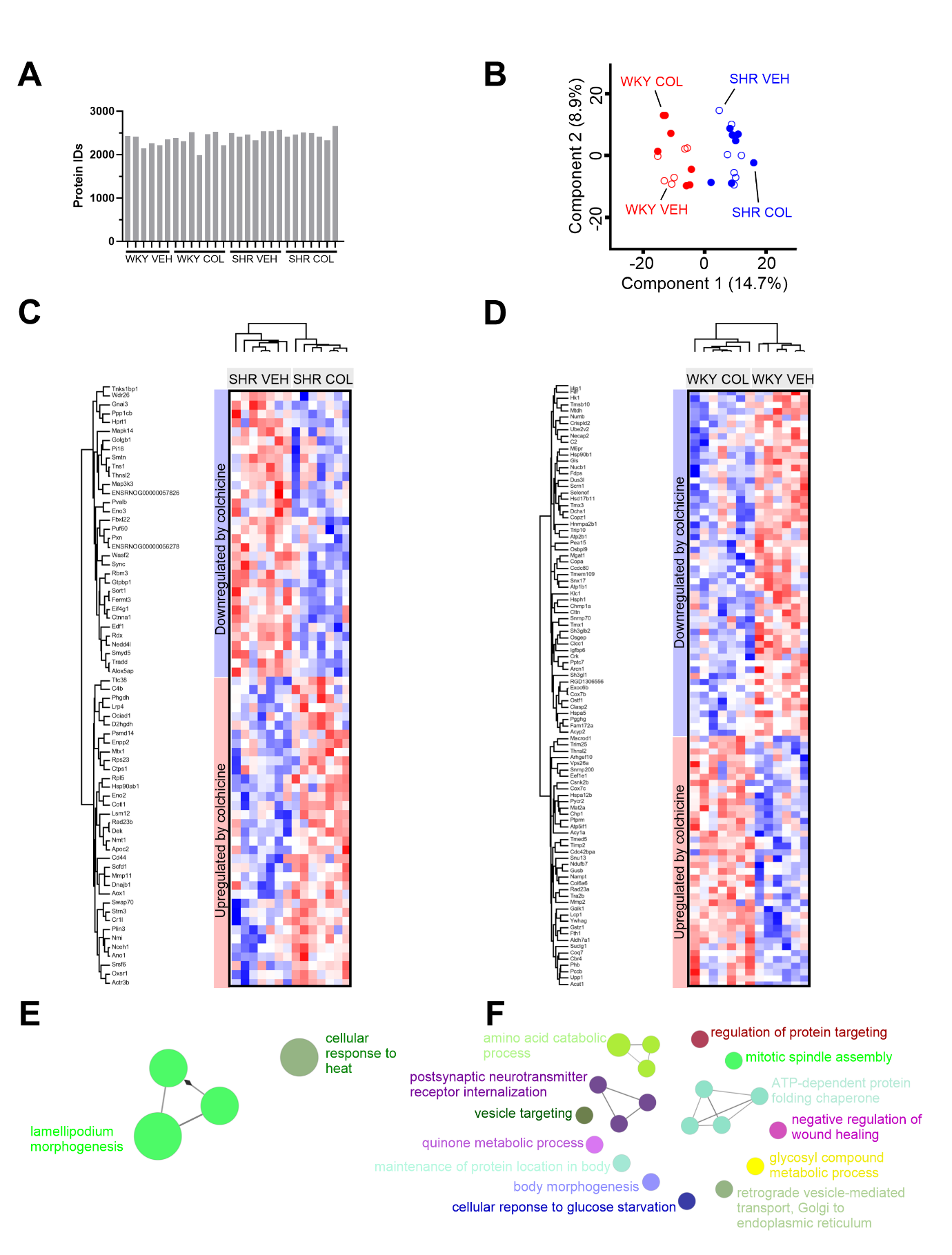

***Supplementary figure S3: Protein profiling of colchicine treatment in aorta.***

*(A) Histogram of unique proteins identified by DIA-MS at 1% false-discovery rate (FDR) identified in spontaneously hypertensive rats (SHR) or normotensive Wistar Kyoto (WKY) controls treated with vehicle (VEH) or colchicine (COL).*

*(B) Principal component analysis (PCA) plot of log2-transformed intensities associated with the samples. Components 1 and 2 are presented.*

*(C & D) Unsupervised hierarchical clustering of significantly regulated proteins identified in spontaneously hypertensive rats (SHR) or normotensive Wistar Kyoto (WKY) controls treated with vehicle (VEH) or colchicine (COL). z-scored values are depicted (-2.5 ‘blue’ to 2.5 ‘red’).*

*(E & F) Gene enrichment analysis of differentially expressed proteins (from C and D) using the GO Biological Process database.*

*Figure S4*

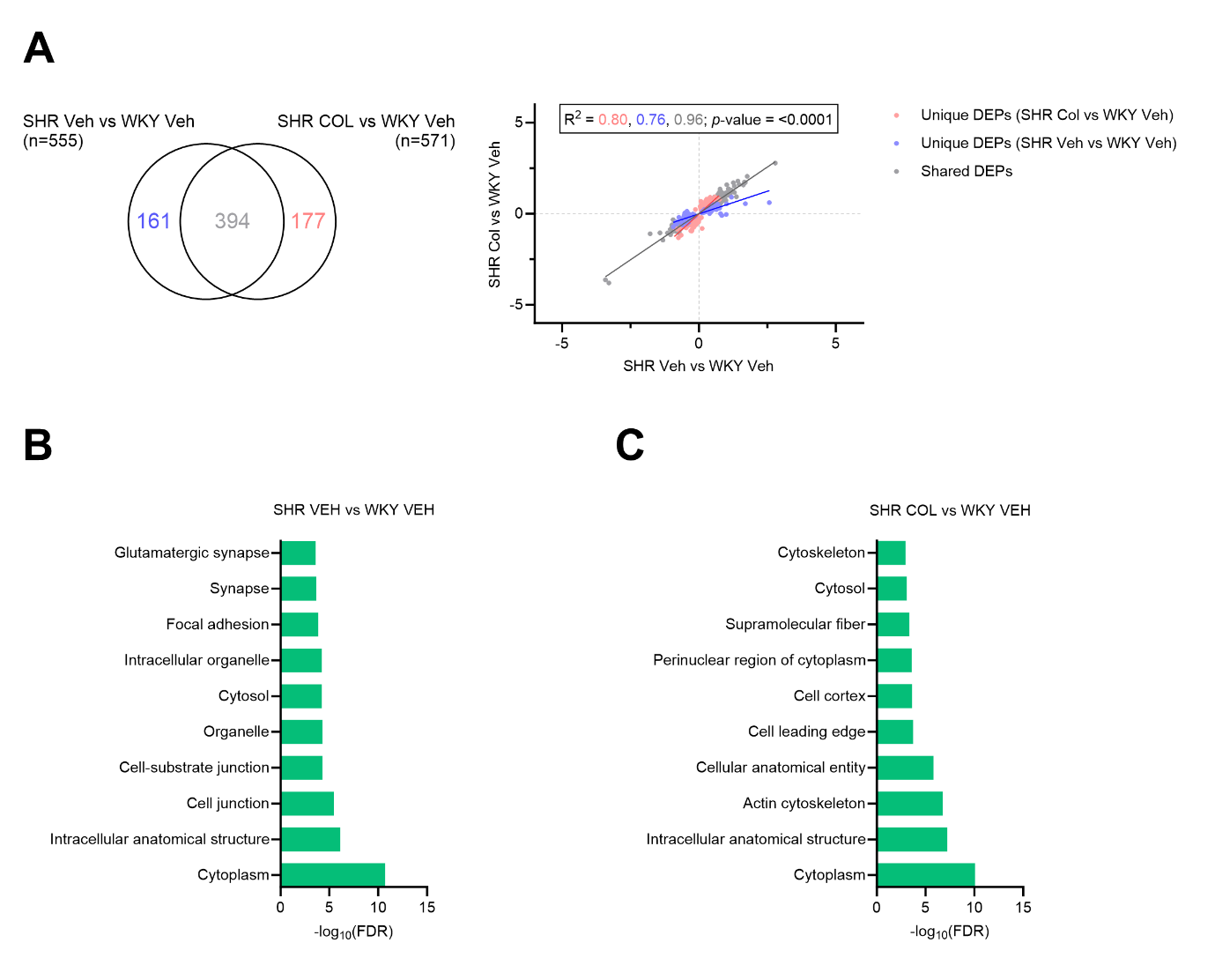

***Supplementary figure S4: Effect of colchicine treatment in aorta.***

*(A) Venn diagram showing unique and shared differentially expressed proteins (DEPs) from two comparisons; Wistar Kyoto (WKY) vehicle (VEH) vs Spontaneously Hypertensive Rat (SHR) vehicle and WKY vehicle vs SHR colchicine (COL). The changes in the expression of proteins in these comparisons is shown in a scatter plot, with R^2^ values calculated. Unique to WKY vehicle vs SHR vehicle = blue, unique to WKY vehicle vs SHR colchicine = red, shared = grey.*

*(B & C) Significance score identified in enrichment analysis of uniquely regulated proteins* *(top10 pathways based significance are shown). Pathway analysis is based on the GO Cellular Component database. Log10 transformed significance score is depicted.*

*Figure S5*

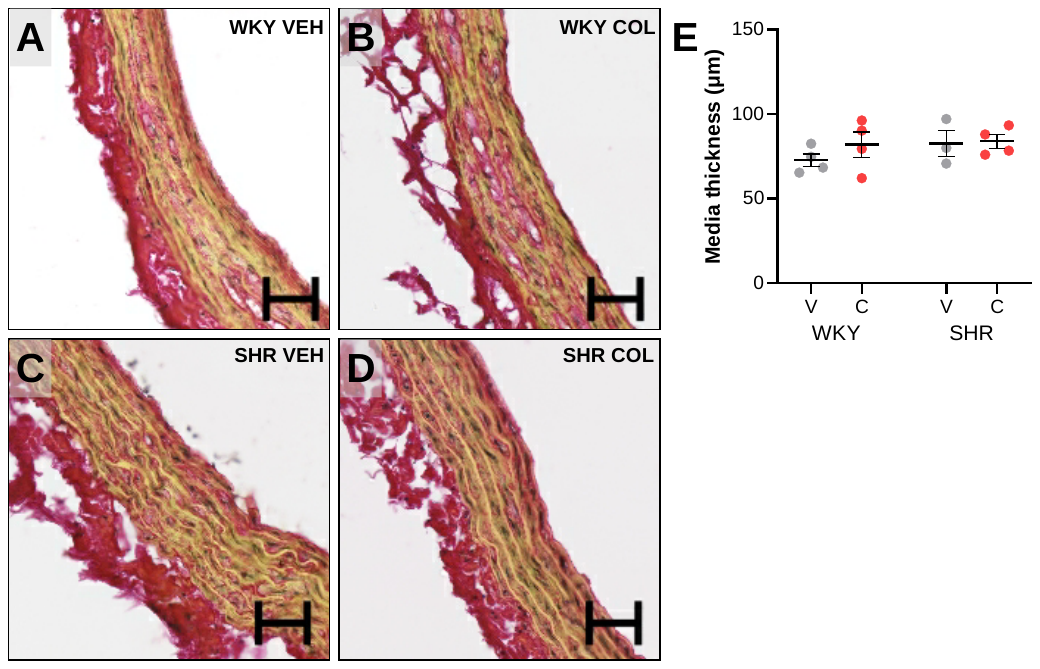

***Supplementary figure S5: Media thickness analysis in thoracic aorta segment****.*

*(A-D) Representative Sirius red stained aorta from the four groups (Wistar Kyoto (WKY) and spontaneously hypertensive rat (SHR) receiving either colchicine (COL) or vehicle (VEH)). Scale bar represents 50 μm; 20 × lens.*

*(E) Mean data ± SEM showing the media thickness in WKY and SHR groups receiving vehicle or colchicine (n = 4 in each group). Statistical significance was assessed by a 2-way ANOVA.*

*Figure S6*

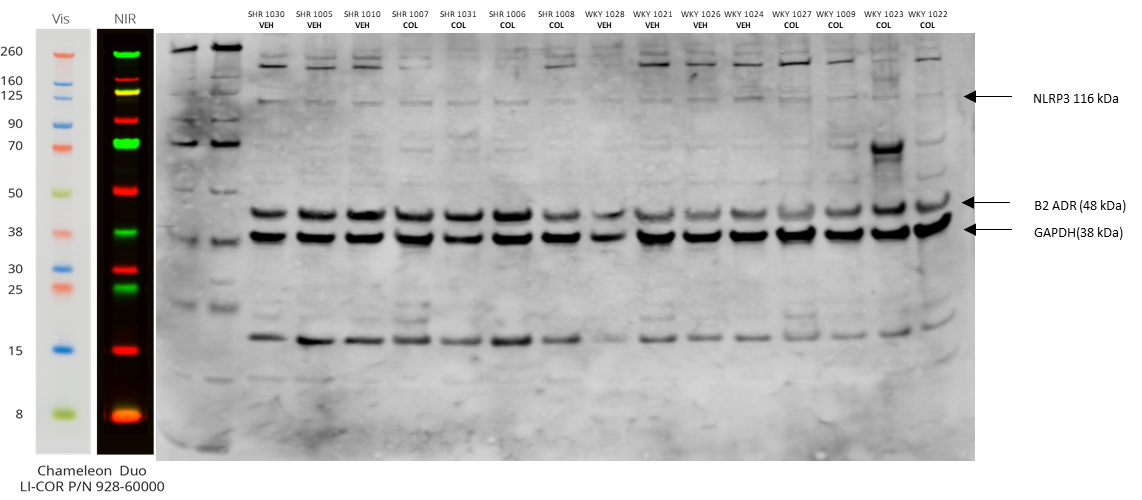

***Supplementary figure S6: Full Western blot of NLRP3 expression in protein lysates from Wistar Kyoto and hypertensive (SHR) rats.***

*NLRP3 expression in 3rd order rat mesenteric arteries from Wistar Kyoto rats (WKY) and Spontaneously Hypertensive Rats (SHR) treated with vehicle (VEH) or colchicine (COL). NLRP3 expression was normalised to GAPDH. The β2 adrenoceptor was also stained for on this blot, as indicated (B2 ADR), but not used for this study. Ladder is shown on the left according to the manufacturer and corresponds to the first 2 lanes of the blot.*

*Figure S7*

A

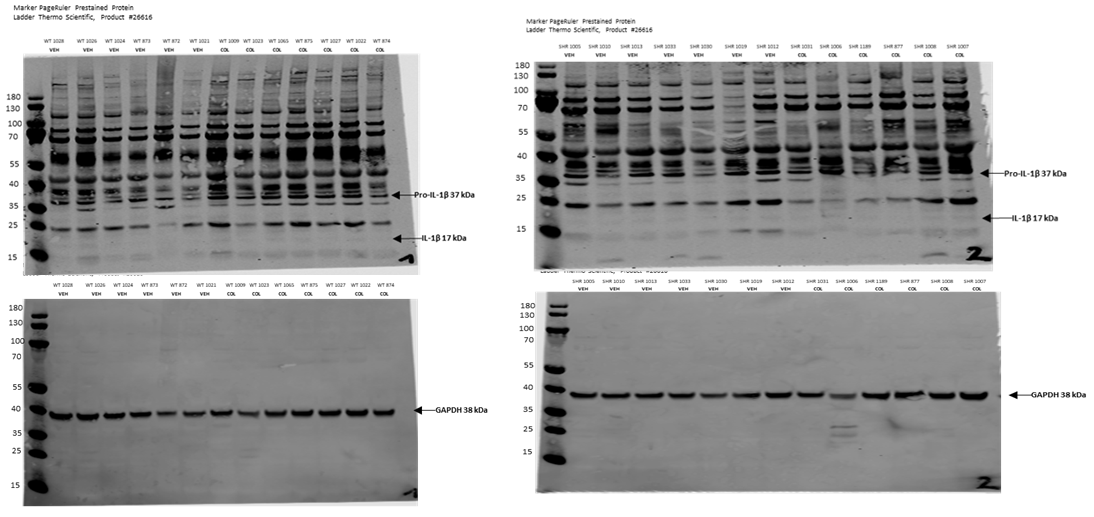

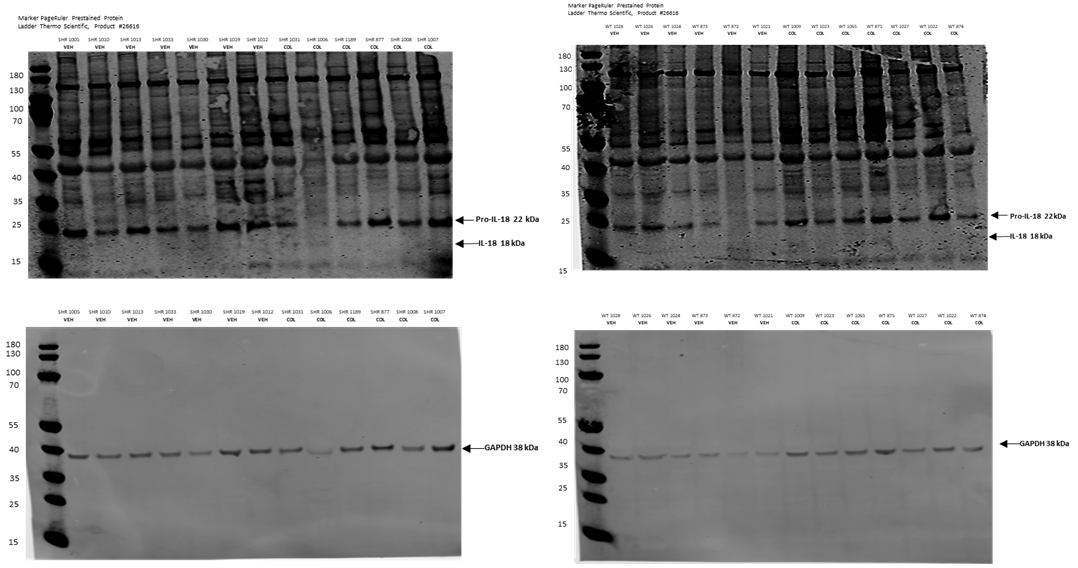

B

***Supplementary figure S7: Representative Western blot of pro and cleaved IL-1β and IL-18 expression in protein lysates from Wistar Kyoto and hypertensive (SHR) rats.***

*(A) Pro and cleaved IL-1β expression in 3rd order rat mesenteric arteries from Wistar Kyoto rats (WKY) and Spontaneously Hypertensive Rats (SHR) treated with vehicle (VEH) or colchicine (COL). IL-1β expression was normalised to GAPDH. Ladder is shown in the left lanes and is marked according to the manufacturer.*

*(B) Pro IL-18 expression in 3rd order rat mesenteric arteries WKYs and SHRs treated with vehicle (VEH) or colchicine (COL). We could not reliably detect IL-18 (18 kDa) expression in these samples. Pro-IL-18 expression was normalised to GAPDH. Ladder is shown in the left lanes and is marked according to the manufacturer.*

*Figure S8*

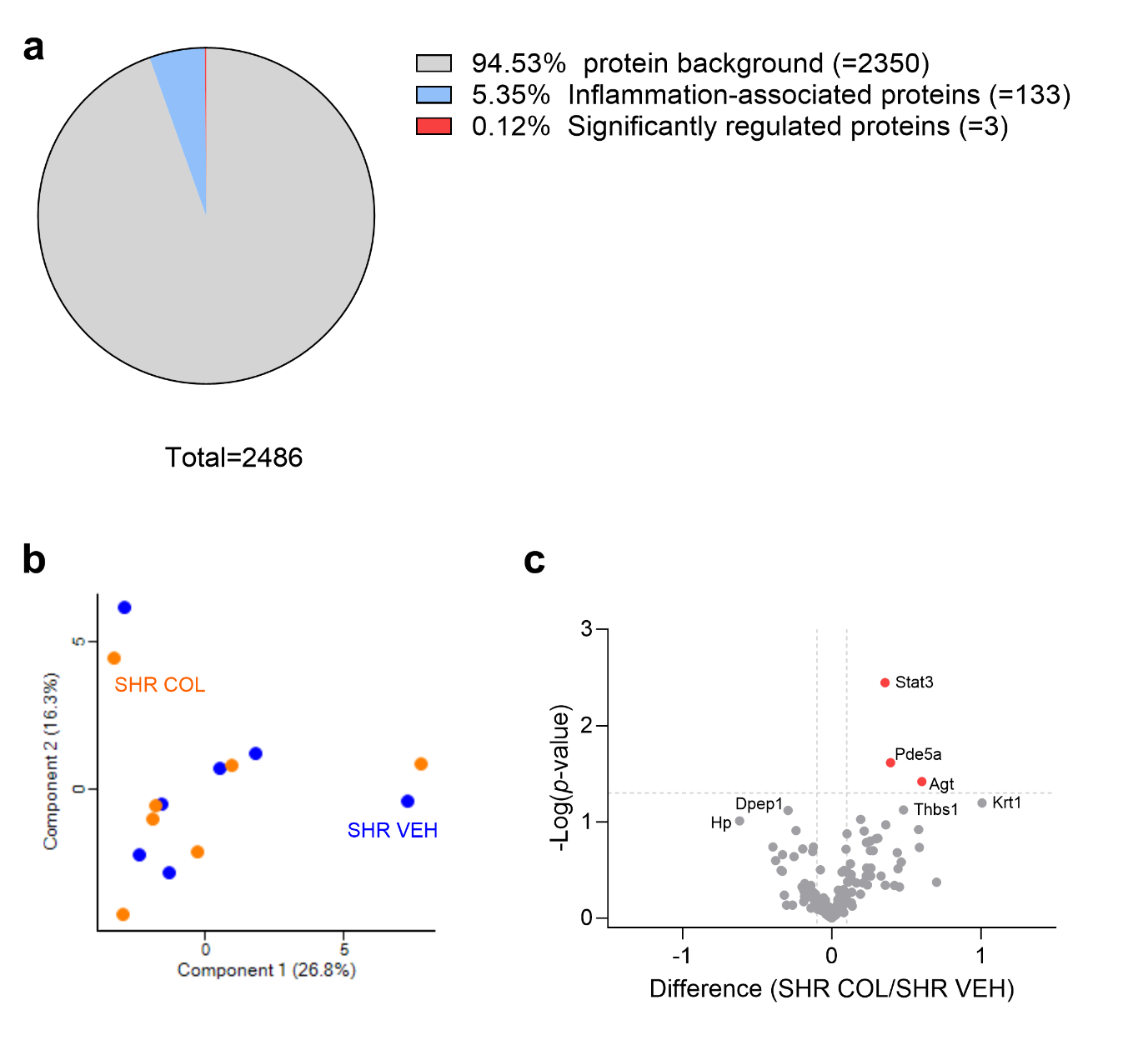
 ***Supplementary figure S8: Inflammation-enrichment analysis in mesenteric artery proteomic data.***

*(A) Pie chart depicting number of proteins in background (grey), inflammation-associated based on inflammation database (blue), and significantly regulated inflammation-associated proteins (red).*

*(B)* *Principal component analysis of spontaneously hypertensive rats (SHR) treated with colchicine (SHR COL; orange) or vehicle (SHR VEH; blue).*

*(C) Volcano plot comparing abundance of inflammation-associated proteins identified in the SHR COL versus SHR VEH comparison. Log2 fold-change differences and non-adjusted p-values are presented. Red = significantly upregulated in SHR COL compared to SHR VEH.*

*Figure S9*

| 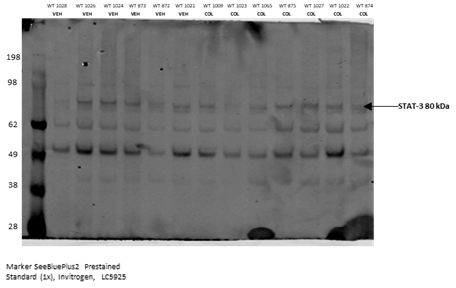 | 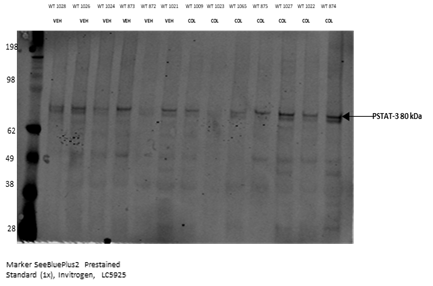 |
| --- | --- |
| 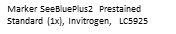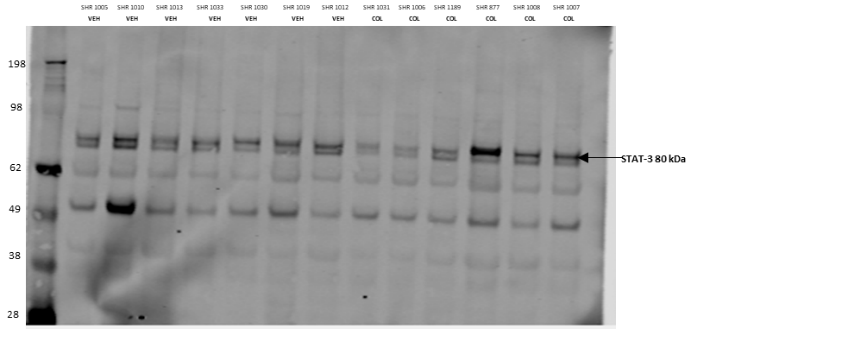 | 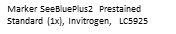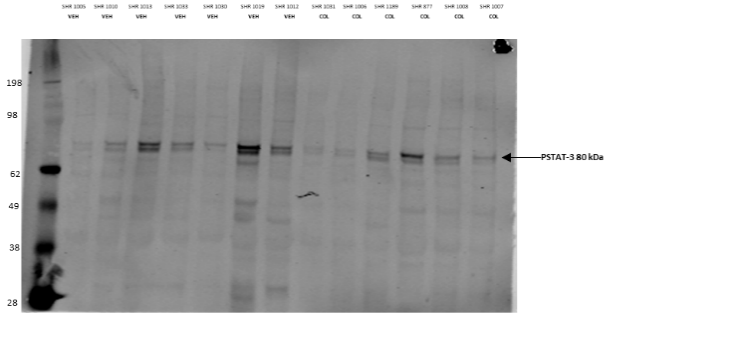 |

***Supplementary figure S9: Representative Western blot of total and phosphorylated STAT3 expression in protein lysates from Wistar Kyoto and hypertensive (SHR) rats.***

*Total (left panel) and phosphorylated (right panel) STAT3 expression in 3rd order rat mesenteric arteries from Wistar Kyoto rats (WKY) and Spontaneously Hypertensive Rats (SHR) treated with vehicle (VEH) or colchicine (COL). Ladder is shown in the left lanes and is marked according to the manufacturer.*

*Figure S10*
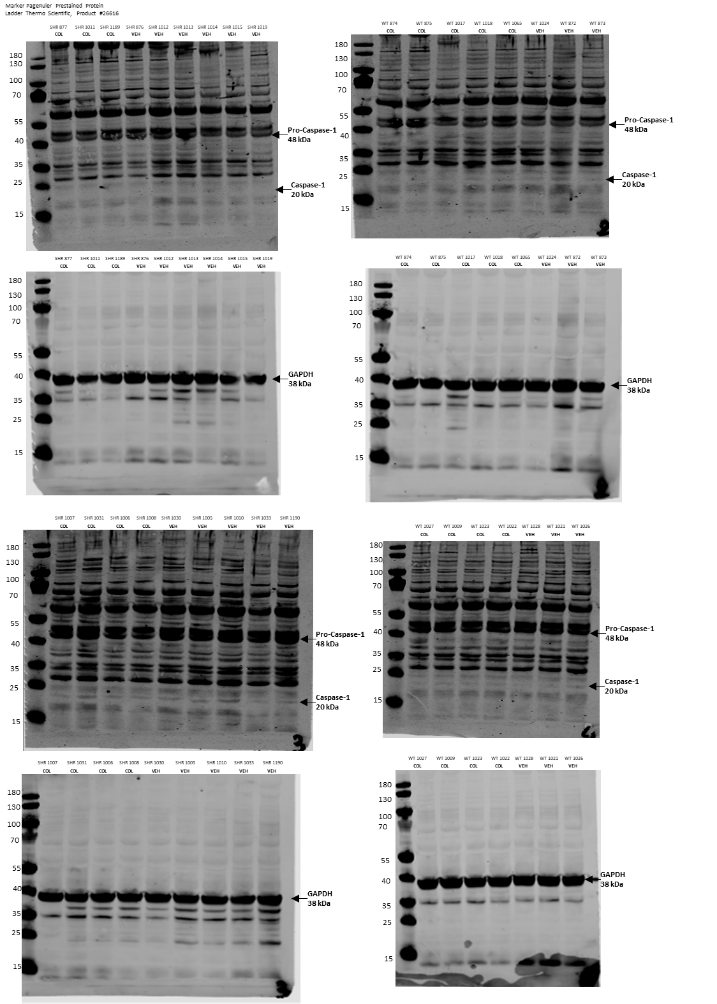

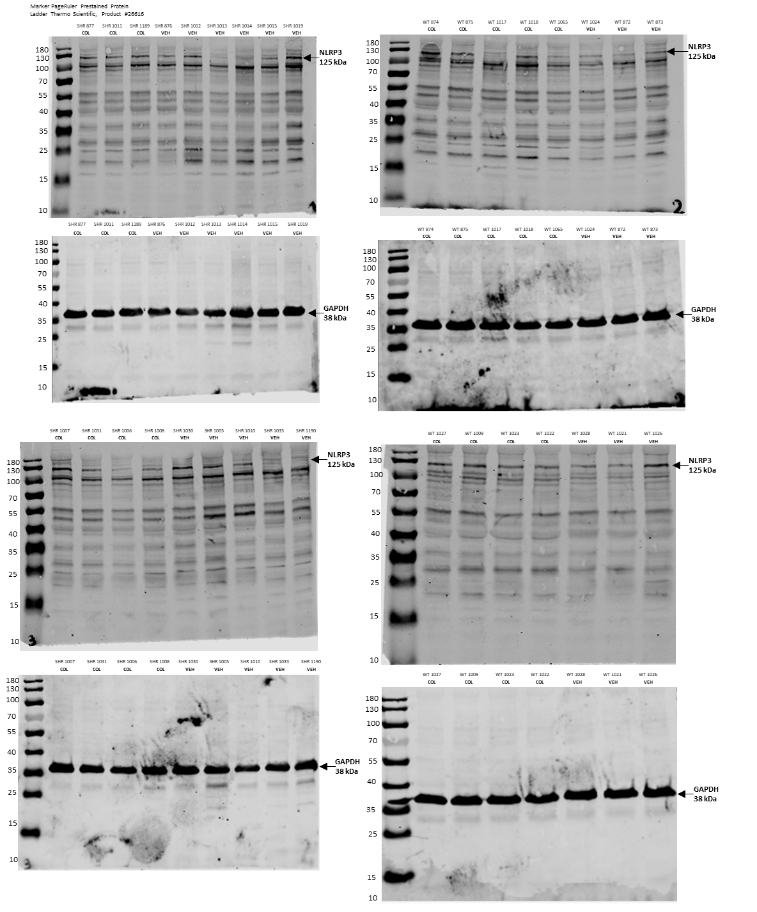

B

A

C

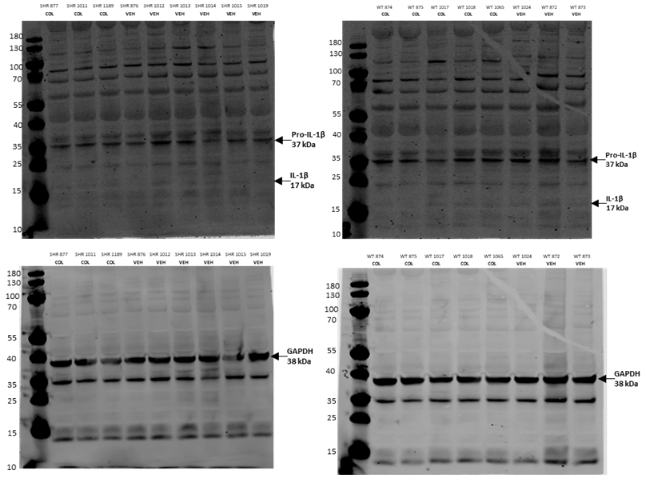

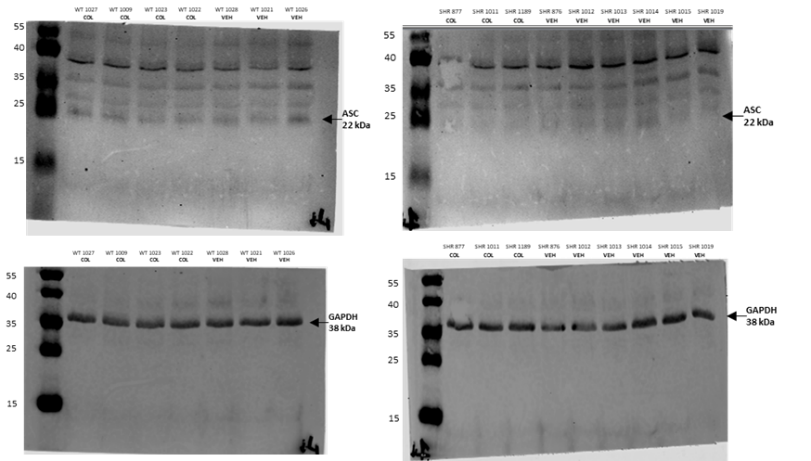

D

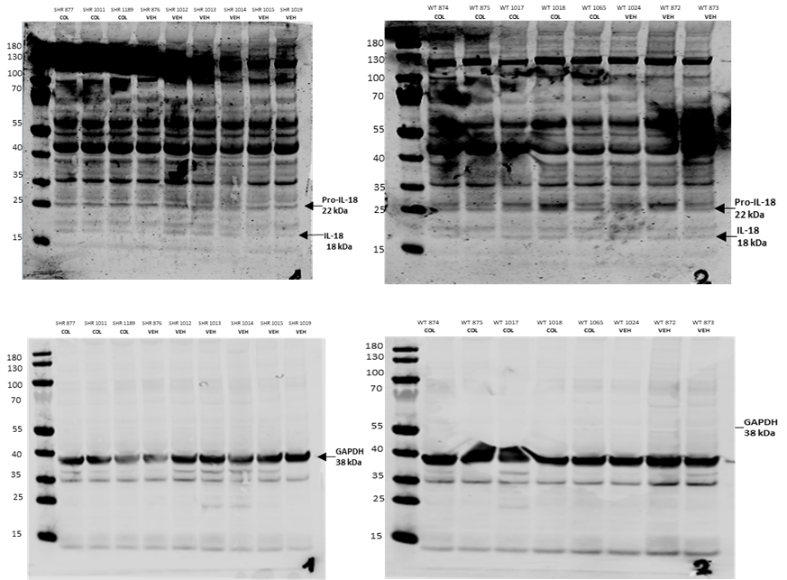

E

***Supplementary figure S10: Representative Western blots of inflammasome-associated proteins in left ventricle lysates from Wistar Kyoto (WKY) and hypertensive (SHR) rats.***

*(A) NLRP3, (B) Pro-caspase and Caspase-1, (C) ASC, (D) Pro-IL-1β and cleaved IL-1β, and (E) Pro-IL-18 and IL-18 expression in the left ventricle of hearts isolated from WKY and SHRs treated with vehicle (VEH) or colchicine (COL). GAPDH expression is also shown for each blot. Ladder is shown in the left lanes and is marked according to the manufacturer.*

***Supplementary Tables***

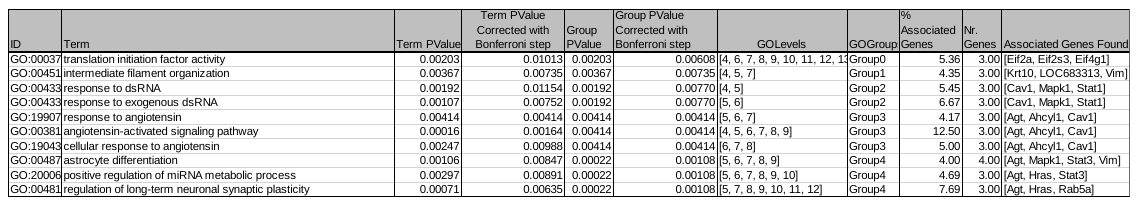
*Table S1*

***Supplementary Table S1: List of Go Terms and associated proteins that are significantly regulated between Spontaneously Hypertensive Rats (SHRs) receiving vehicle vs colchicine.***

*Details of the GO terms (determined from database: GO_BiologicalProcess-EBI-UniProt-GOA-ACAP-ARAP_10.01.2024_00h00) associated with differently regulated proteins between SHRs receiving vehicle compared to colchicine.*

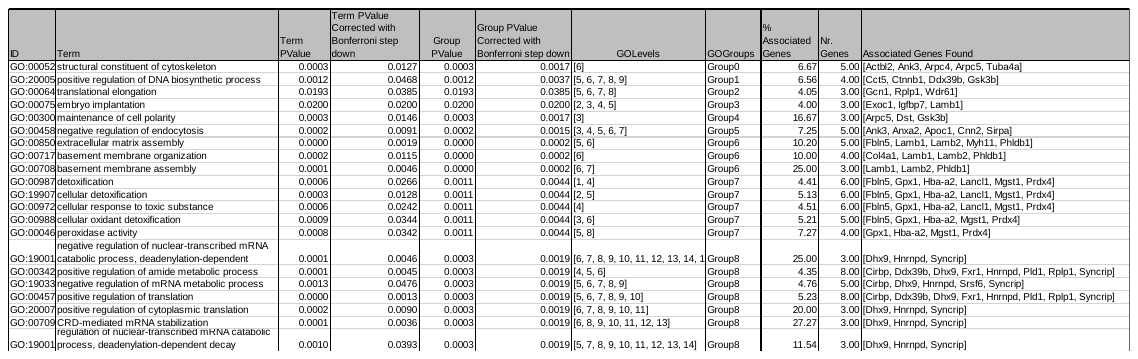
*Table S2*

***Supplementary Table S2: List of Go Terms and associated proteins that are significantly regulated between Wistar Kyoto Rats (WKY) receiving vehicle vs colchicine.***

*Details of the GO terms (determined from database: GO_BiologicalProcess-EBI-UniProt-GOA-ACAP-ARAP_10.01.2024_00h00) associated with differently regulated proteins between WKY rats receiving vehicle compared to colchicine.*

*Table S3*

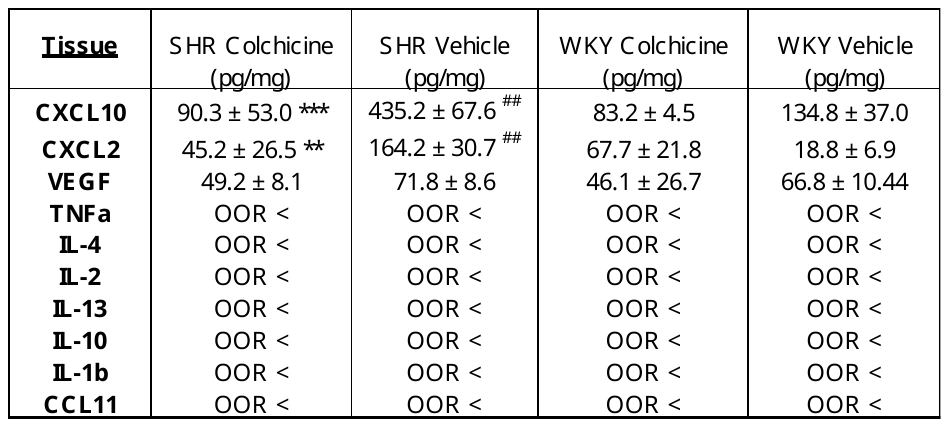

***Supplementary Table S3: Multiplex immunoassay results showing the concentration of several inflammatory markers in mesenteric artery tissue of the spontaneously hypertensive rats (SHR) and Wistar Kyoto rats (WKY) treated with either vehicle or colchicine.***

*Only CXCL10, CXCL2, and VEGF were consistently detected in our samples, whereas TNFα, IL-2, IL-4, IL-10, IL-13, IL-1β and CCL11 were under the detectable range (OOR <) for the assay.* *Statistical significance was determined by 2-way ANOVA followed by a Fisher’s uncorrected LSD; **(p<0.01) and ***(p<0.001) denote significance between SHR colchicine vs SHR Vehicle, ^##^(p<0.01) denotes significance between SHR vehicle and WKY vehicle.*
